# Diversity and Distribution of a Novel Genus of Hyperthermophilic *Aquificae* Viruses Encoding a Proof-reading Family-A DNA Polymerase

**DOI:** 10.1101/2020.02.27.968263

**Authors:** Marike Palmer, Brian P. Hedlund, Simon Roux, Philippos K. Tsourkas, Ryan K. Doss, Casey Stamereilers, Astha Mehta, Jeremy A. Dodsworth, Michael Lodes, Scott Monsma, Tijana Glavina del Rio, Thomas W. Schoenfeld, Emiley A. Eloe-Fadrosh, David A. Mead

## Abstract

Despite the high abundance of *Aquificae* in many geothermal systems, these bacteria are difficult to culture and no viruses infecting members of this phylum have been isolated. Here, we describe the complete, circular dsDNA Uncultivated Virus Genome (UViG) of *Thermocrinis* Octopus Spring virus (TOSV), derived from metagenomic data, along with eight related UViGs representing three additional species, *Thermocrinis* Great Boiling Spring virus (TGBSV), *Aquificae* Joseph’s Coat Spring Virus (AJCSV), and *Aquificae* Conch Spring Virus (ACSV). Four near-complete UViGs, ranged from 37,256 bp to 41,208 bp and encoded 48 to 53 open reading frames. Despite low overall similarity between viruses from different hot springs, the genomes shared a high degree of synteny, and encoded numerous genes for nucleotide metabolism, including a polyprotein PolA-type polymerase with likely accessory functions, a DNA Pol III beta subunit (sliding clamp), a thymidylate kinase, a DNA gyrase, a helicase, and a DNA methylase. Also present were conserved genes predicted to code for phage capsids, large and small terminases, portal protein, holin, and lytic transglycosylase, all consistent with a distant relatedness to cultivated *Caudovirales*. TOSV and TGBSV had the highest coverage in their respective metagenomes and are predicted to infect *Thermocrinis ruber* and *Thermocrinis jamiesonii*, respectively, as multiple CRISPR spacers matching the viral genomes were identified within *Thermocrinis ruber* OC1/4^T^ and *Thermocrinis jamiesonii* GBS1^T^. Based on the predicted, unusual bi-directional replication strategy, low sequence similarity to known viral genomes, and a unique position in gene-sharing networks, we propose a new putative genus, Pyrovirus, in the order *Caudovirales*.

## INTRODUCTION

Viruses are the most abundant biological entities on Earth and are important drivers of genetic exchange, secondary production, and host metabolism on both local and global scales (Breitbart et al., 2018; Fuhrman 1999; Rohwer and Thurber 2009; Suttle 2007). They also possess a high density of nucleic acid-synthesis and -modifying enzymes that are important sources of enzymes for the biotechnology sector. Despite their importance, cultivation of viruses in the laboratory is limited by challenges associated with cultivating their hosts. This problem is particularly true for viruses of thermophiles and hyperthermophiles because many hosts remain uncultured. Also, most thermophiles do not readily form lawns on solid media, which are typically exploited to screen for plaques. Although direct observation of filtrates from geothermal springs and enrichments has revealed a high diversity of virus morphotypes (Rice et al., 2001; Rachel et al., 2002), few thermophilic viruses have been studied in enrichment cultures and even fewer have been isolated in culture with their host. Currently, the NCBI Viral Genomes database lists 59 thermophilic archaeal viruses out of 95 total genomes, representing ten families (http://www.ncbi.nlm.nih.gov/genomes/GenomesGroup.cgi?taxid=10239&host=archaea; accessed 2/3/20); however, 49 of these infect members of the thermoacidophilic family *Sulfulobaceae*, leaving other archaeal thermophiles vastly under-explored. Similarly, only 15 of the 2,500 bacteriophage genomes represent thermophilic or hyperthermophilic viruses, representing only three virus families (http://www.ncbi.nlm.nih.gov/genomes/GenomesGroup.cgi?taxid=10239&host=bacteria; accessed 2/3/20). Strikingly, although members of the phylum *Aquificae* (syn. *Aquificota*) predominate in many terrestrial and marine high-temperature ecosystems (Reysenbach et al., 2005; Spear et al., 2005), to date, no cultivated viruses infecting *Aquificae* have been described.

Microbial ecologists have increasingly turned to cultivation-independent approaches to probe microbial diversity in nature. Although the low nucleic acid content and lack of universal conserved marker genes slowed the development of viral metagenomics, this field is now in full swing (Emerson et al., 2018; Koonin et al., 2018; Paez-Espino et al., 2016). One of the early viral metagenomic investigations focused on Octopus Spring and other circumneutral pH springs in Yellowstone National Park (Schoenfeld et al., 2008), revealing 59 putative DNA polymerase (*pol*) genes, which were subsequently screened for heterologous activity in *E. coli* (Moser et al., 2012). The most thermophilic of these enzymes, 3173 PolA, also demonstrated high-fidelity, thermostable reverse-transcriptase (RT) activity, and strand-displacement activity and was subsequently marketed by Lucigen Corporation as a single-enzyme RT-PCR system called PyroPhage and RapidDxFire. That enzyme was further improved by molecular evolution and fusion of a high-performance chimeric variant of 3173 PolA with the 5’ to 3’ exonuclease domain of *Taq* polymerase to improve probe-based detection chemistries and enable highly sensitive detection of RNA (Heller et al., 2019).

A study of the diversity and evolution of 3173 PolA and related polymerases revealed clues about its complex evolutionary history (Schoenfeld et al., 2013). In addition to their discovery in viral metagenomes from hot springs, 3173 *polA*-like genes were also detected in two of the three families of *Aquificae*, where they have orthologously replaced host DNA *polA* genes, and phylogenetically diverse, non-thermophilic bacteria, where they appear to be transient alternative *polA* genes, presumably due to recombination following non-productive infections. Amazingly, 3173 *polA*-like genes are also known to encode thermophilic, nuclear-encoded, apicoplast-targeted polymerases in eukaryotic parasites in the *Apicomplexa* (e.g., *Plasmodium*, *Babesia*, and *Toxoplasma*) (Seow et al., 2005). The origin of these genes likely involved fixation of a progenitor sequence into the nuclear genome following endosymbiosis of a red alga (proto-apicoplast) containing a bacterial symbiont carrying a viral *polA* (Schoenfeld et al., 2013).

Recently, an Uncultivated Virus Genome (UViG) containing the 3173 *polA* gene was described (Mead et al., 2018). Here, we further describe the OS3173 virus genome and related UViGs, including nearly complete UViGs from several Yellowstone springs and Great Boiling Spring (GBS), Nevada, that range from 37,256 bp to 41,208 bp and encode 48 to 53 open reading frames. The presence of fragments of these genomes in CRISPR arrays encoded by *Thermocrinis ruber* OC1/4^T^, *Thermocrinis jamiesonii* GBS1^T^, *Hydrogenobaculum* sp. 3684, and *Sulfurihydrogenibium yellowstonense* SS-5 ^T^ genomes, along with similarity between many viral genes and *Aquificaceae* genes, supports the previous hypothesis (Mead et al., 2018; Schoenfeld et al., 2013) that *Thermocrinis* and probably other *Aquificae* are putative hosts of these viruses. The high abundance of these viruses and their hosts suggests they may play an important role in chemolithotrophic productivity in geothermal springs globally, in addition to their role in evolution as a vector for horizontal gene transfer.

## RESULTS AND DISCUSSION

### DOMINANT VIRAL UViGS FROM OCTOPUS SPRING AND GREAT BOILING SPRING ENCODES AN UNUSUAL DNA POLYMERASE

Viral particles were isolated from Octopus Spring in Yellowstone National Park and Great Boiling Spring (GBS) in the U.S. Great Basin by sequential tangential-flow filtration (Schoenfeld et al., 2008) and used for metagenomic sequencing (Mead et al., 2018). In parallel, the cell fraction from GBS was also used for metagenomic sequencing. Forty-three percent of the reads from the Octopus Spring virus-enriched metagenome assembled into a single contig herein called *Thermocrinis* Octopus Spring virus (TOSV, equivalent to the term OS3173 used previously (Mead et al., 2018)). The TOSV genome was 37,256 bp and encoded 48 predicted open reading frames (Figure 1A, S1A), as detailed below. Metagenomic coverage was high (mean 913X) and uniform across the TOSV genome above 95% nucleotide identity, and read depth was low at lower identity (Figure S1B,C). Together, these data indicate that TOSV was likely the dominant virus present at the time and place of sampling. Among the 48 predicted genes (Supplementary File S3) was a full-length, polyprotein PolA-type polymerase nearly identical to 3173 PolA (Figure 2), a portion of which was previously discovered via Sanger sequencing of metagenomic clone libraries (Schoenfeld et al., 2008). The near-complete absence of TOSV reads from a pink streamer microbial metagenome dominated by *Thermocrinis* from the outflow of Octopus Spring (Takacs-Vesbach et al., 2013) suggests viral activity is temporally or spatially variable in that environment (Supplementary File S1).

**Figure 1.**
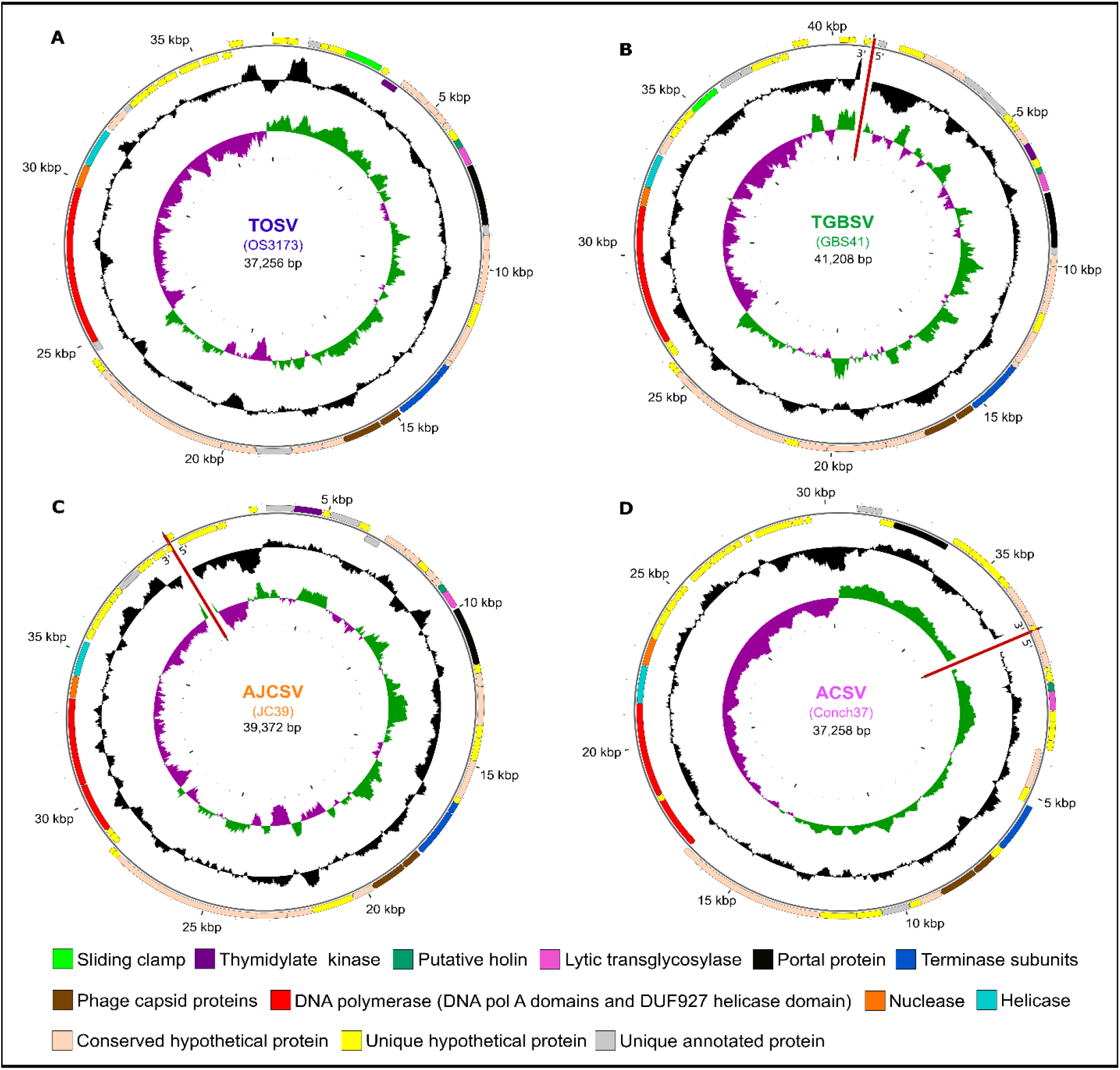
Map of four large UViGs. The uncultivated viral genomes were recovered from metagenomes from (A) Octopus Spring (TOSV); (B) Great Boiling Spring (TGBSV); (C) Joseph’s Coat Spring (AJCSV); and (D) Conch Spring (ACSV). Outer circles show ORFs and selected annotation features, with arrows in the putative direction of transcription. Middle circles show the GC content and the inner circles show the GC skew. The sequences of GBS41, JC39 and Conch37 could not be circularized as indicated with red lines. Maps have been rotated to reflect the orientation of OS3173. TOSV, TGBSV, AJCSV, and ACSV are represented by OS3173, GBS41, JC39, and Conch37, respectively.

**Figure 2.**
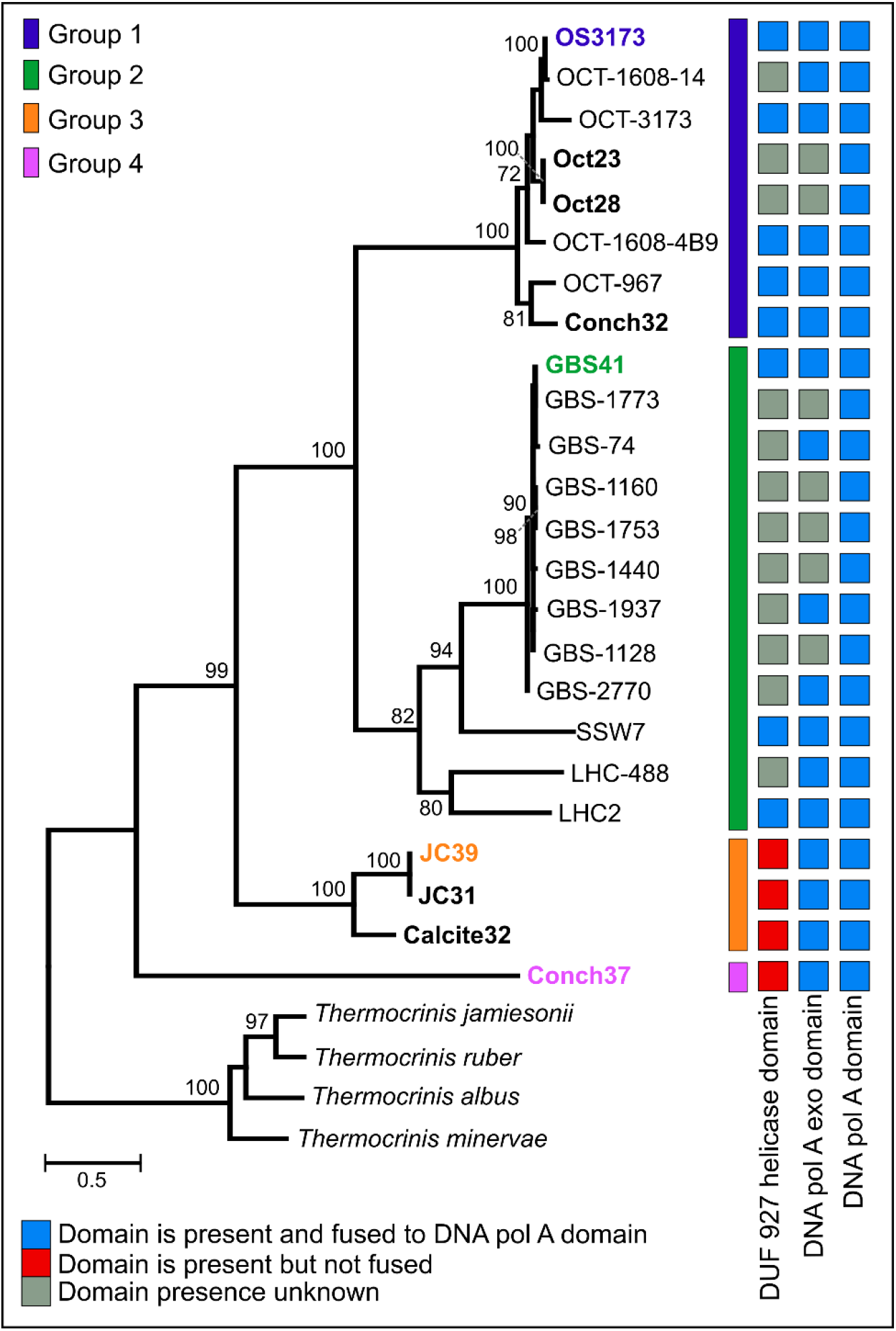
Phylogeny and structure of 3173 PolA-like proteins. Maximum-likelihood phylogeny of near full-length 3173 PolA-like proteins, with bootstrap values above 70% from 1,000 pseudoreplicates indicated. OCT, Oct or OS, Octopus Spring; Conch, Conch Spring; GBS, Great Boiling Spring; SSW, Sandy’s Spring West; LHC, Little Hot Creek; JC, Joseph’s Coat Spring; Calcite, Calcite Spring. The presence of helicase, exonuclease, and polymerase domains are indicated, where known. The scale bar indicates the number of amino acid changes per site. Taxa indicated in bold represent UViGs that were >23kb, while the representative UViG of each group is colored in the corresponding group color. OS3173, GBS41, JC39, and Conch37 represent TOSV, TGBSV, AJCSV, and ACSV, respectively.

Other viral contigs with lower coverage present in the Octopus Spring virus-enriched metagenome (Figure 4, S2, S3) were similar to *Pyrobaculum* Spherical Virus (PSV) (Häring et al., 2004), a member of the *Globuloviridae*, which was previously described in Octopus Spring viral metagenomes (Schoenfeld et al., 2008; Mead et al., 2018), or distantly related to *Siphoviridae* viruses infecting mesophilic *Actinobacteria* or *Leptospira* (Figure S3) (Supplementary File S1).

A similar viral contig encoding a 3173 PolA-like protein (Figure 2), herein putatively named *Thermocrinis* Great Boiling Spring virus (TGBSV), was obtained from the GBS cell metagenome. The TGBSV genome is 41,208 bp and encodes 53 putative open reading frames (Figure 1B, S4A). Genomic coverage was low across the majority of the genome (mean 15.4X), yet it was highly variable in the intergenic regions on either end of the linear contig (Figure S4B,C). TGBSV reads were also recruited from the GBS virus-enriched metagenome at 50.4X coverage, where TGBSV was the viral contig with the highest coverage (Supplementary File S1), although the *de novo* assembly was fragmented. In contrast to Octopus Spring, the high recruitment of viral reads from the GBS cellular metagenomes suggests active infection of *Thermocrinis jamiesonii* in GBS during the time of sampling.

Other contigs from the GBS virus-enriched metagenome (Figure S5, S6) were distantly related to viruses from halophilic *Euryarchaeota*, various *Sulfolobales* viruses, and PSV (Figure S6). *Pyrobaculum* is relatively abundant in GBS (Costa et al., 2009; Cole et al., 2013); however, *Sulfolobales* are not known to occur at GBS, as no high-temperature, low-pH habitat is known to exist there. Due to the small size of these contigs and large genetic distance to characterized relatives, these relationships are highly uncertain.

The virus-enriched metagenomes from Octopus Spring and GBS are summarized in Supplementary File S1, including read recruitment, vContact 2.0 files, and CRISPR spacer matches of the 10 viral contigs with the highest coverage from these metagenomes.

### RECOVERY OF TOSV-LIKE GENOMES FROM YELLOWSTONE AND GREAT BASIN SPRING METAGENOMES

To assess the distribution and diversity of similar viruses, the full-length 3173 PolA gene of TOSV was used to recruit homologs *in silico* from public databases. In total, 23 unique contigs containing 3173 *polA*-like genes were obtained from cell and virus-enriched metagenomes from Yellowstone and U.S. Great Basin hot springs (Table 1) (Figure 2). The Yellowstone springs, specifically Octopus Spring, Conch Spring, Joseph’s Coat Spring (Scorodite Spring), and Calcite Spring, span several geothermal areas; each is circumneutral (pH 6.0 to 8.8) and has a source that is boiling or near-boiling, and several are known to host abundant populations of *Aquificae* (Reysenbach et al., 1994; Reysenbach et al., 2000). In the Great Basin, Great Boiling Spring and Sandy’s Spring West are only ~1 km apart (Costa et al., 2009), but Little Hot Creek is ~380 km away, and each is separated from the Yellowstone springs by >1,200 km. These springs also share a circumneutral pH, near-boiling sources, and abundant *Aquificae* populations (Costa et al., 2009; Cole et al., 2013; Vick et al., 2010).

**TABLE 1.**
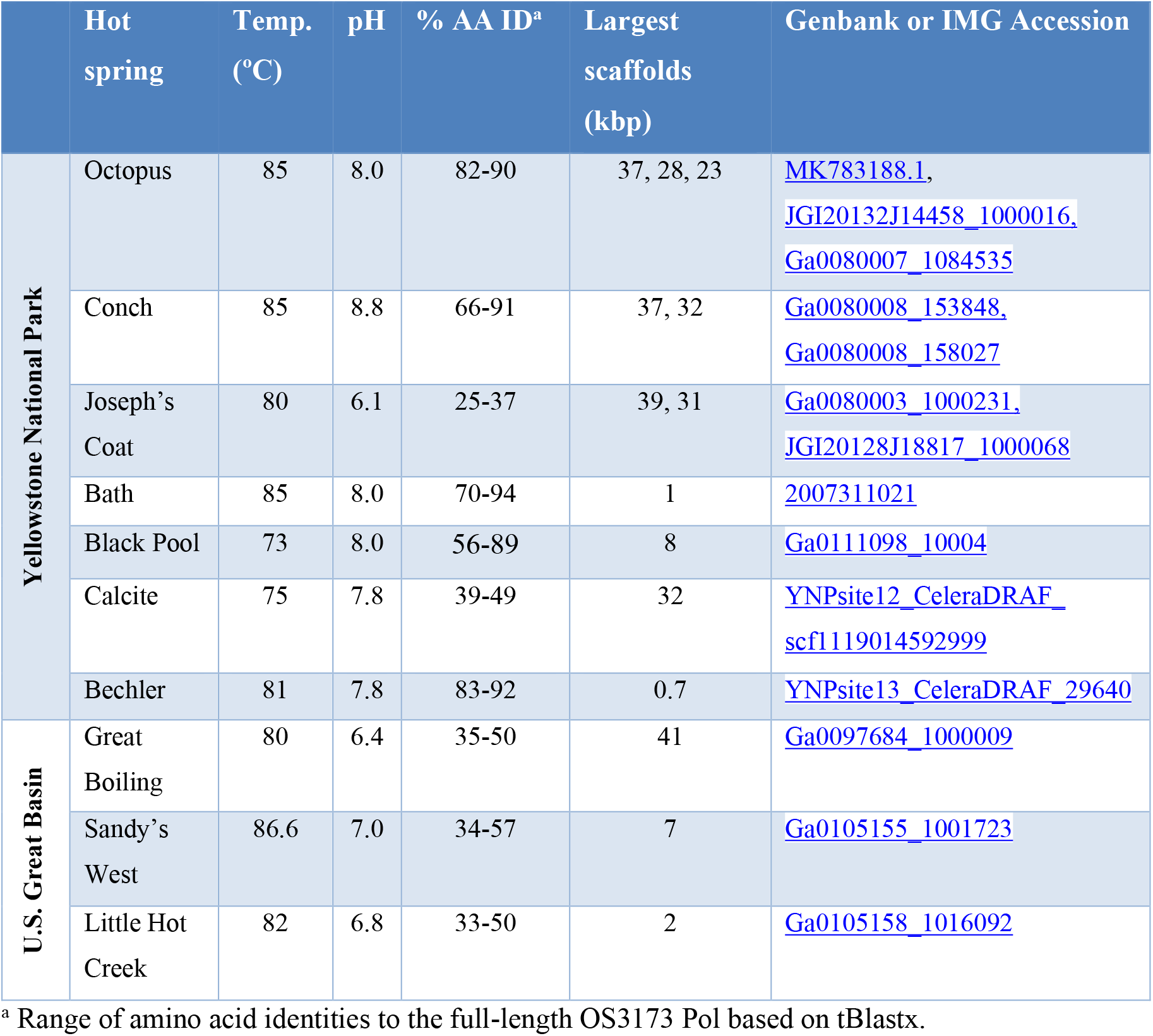
Distribution of OS3173-like *polA* genes in metagenomic databases.

|  | Hot<br>spring | Temp.<br>(°C) | pH | % AA ID<sup>a</sup> | Largest<br>scaffolds<br>(kbp) | Genbank or IMG Accession |
| --- | --- | --- | --- | --- | --- | --- |
| Yellowstone National Park | Octopus | 85 | 8.0 | 82-90 | 37, 28, 23 | MK783188.1,<br>JGI20132J14458_1000016,<br>Ga0080007_1084535 |
|  | Conch | 85 | 8.8 | 66-91 | 37, 32 | Ga0080008_153848,<br>Ga0080008_158027 |
|  | Joseph's<br>Coat | 80 | 6.1 | 25-37 | 39, 31 | Ga0080003_1000231,<br>JGI20128J18817_1000068 |
|  | Bath | 85 | 8.0 | 70-94 | 1 | 2007311021 |
|  | Black Pool | 73 | 8.0 | 56-89 | 8 | Ga0111098_10004 |
|  | Calcite | 75 | 7.8 | 39-49 | 32 | YNPsite12_CeleraDRAF_scf1119014592999 |
|  | Bechler | 81 | 7.8 | 83-92 | 0.7 | YNPsite13_CeleraDRAF_29640 |
| U.S. Great Basin | Great<br>Boiling | 80 | 6.4 | 35-50 | 41 | Ga0097684_1000009 |
|  | Sandy's<br>West | 86.6 | 7.0 | 34-57 | 7 | Ga0105155_1001723 |
|  | Little Hot<br>Creek | 82 | 6.8 | 33-50 | 2 | Ga0105158_1016092 |

<sup>a</sup> Range of amino acid identities to the full-length OS3173 Pol based on tBlastx.

Phylogenetic analysis of the near-complete 3173 PolA-like proteins revealed four well-supported groups that were mostly site-specific (Figure 2), except that one of two Pols from Conch Spring grouped with several from Octopus Spring in Group 1, whereas a distinct Conch Spring Pol split off at the most basal node in the phylogeny (Group 4). Additionally, the Pols from the two pyrite-precipitating springs, Joseph’s Coat Spring and Calcite Spring, grouped together in Group 3. The Pols from Great Basin springs were monophyletic and distinct from the Yellowstone Pols, forming Group 2, following a pattern seen for several thermophilic bacteria and archaea (Dodsworth et al., 2015; Miller-Coleman et al., 2012; Zhou et al., 2019). All the full-length 3173 PolA-like proteins contained a 3’-5’ proofreading exonuclease and DNA polymerase (3’exo/pol) domain, as is typical of many bacterial PolAs. Several also contained putative helicase domains (DUF 927), described later in detail; however, this domain was fused to form a putative polyprotein in Groups 1 and 2, or alternatively present as a separate open reading frame in the four most divergent Pols, all from springs north of Yellowstone Lake (Groups 3 and 4) (Figure 2). Each of the metagenomes contained only one of the Pol variants, except for the previously mentioned Conch Spring Pols.

Nine of the contigs containing the genes encoding the 3173 PolA-like proteins were >23 kbp and were thus considered UViGs (Figure 3, Supplemental Table S1). All nine UViGs were compared by tBlastx to identify other regions of homology and assess genomic synteny (Figure 3). Within the groups previously identified by the Pol phylogeny, shared gene content and synteny were both high. Shared gene content and synteny between the groups was considerably lower, reflecting low average amino acid identities (Figure 4C); however, some of the core genes were organized similarly even in the most distant genomes, including the polymerase/helicase, terminase subunits, and phage capsid proteins, described in detail below.

**Figure 3.**
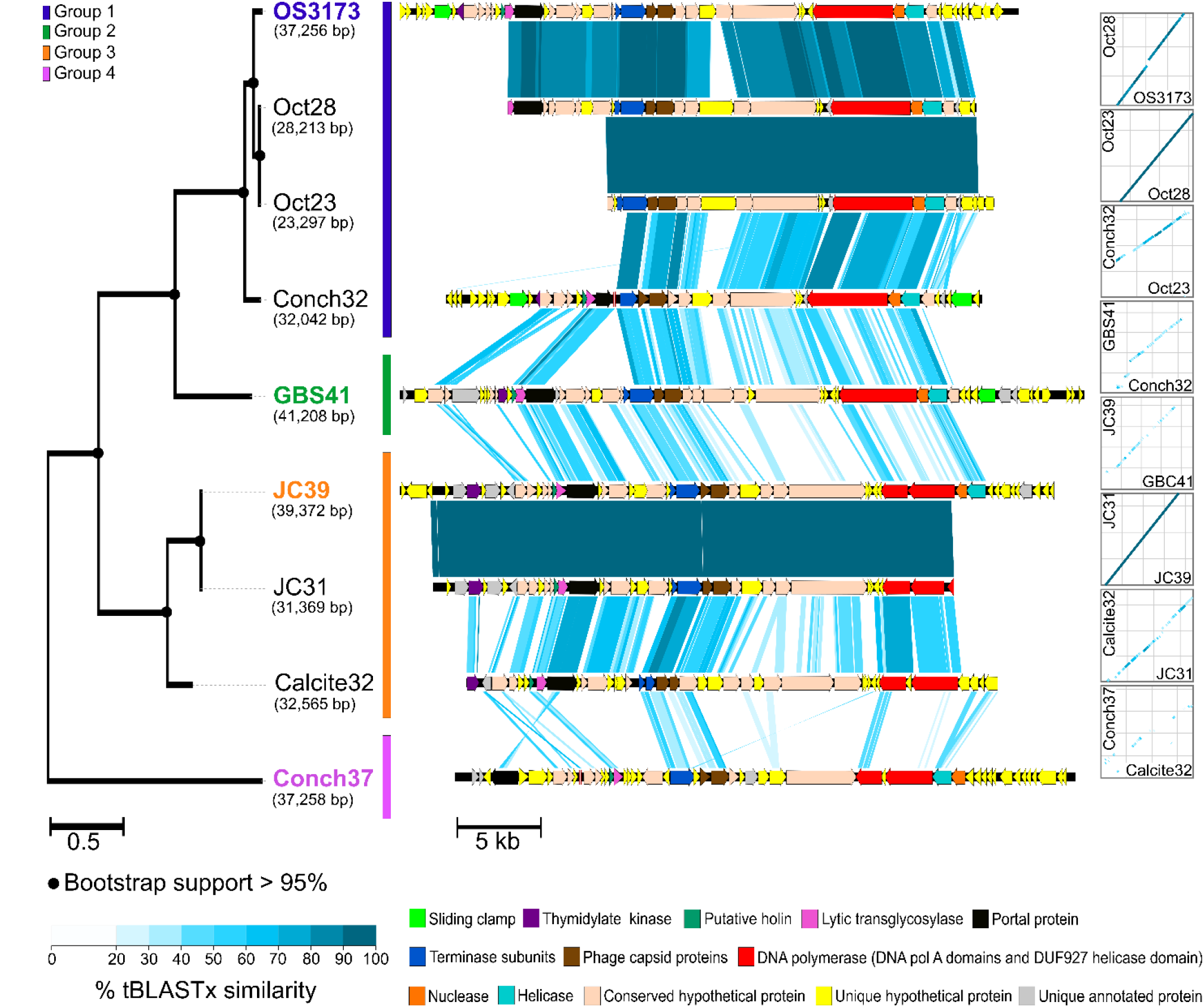
Synteny and amino acid identity across OS3173-like viral genomes. Synteny based on pairwise tBLASTx similarity across OS3173-like viral genomes, showing high overall synteny and few inversions, despite low amino acid identity between groups. The PolA phylogeny for the nine UViGs is indicated on the left of the figure. The representative taxa of each polA-based group are indicated in the group’s corresponding color. Bootstrap support above 95% is indicated at the nodes. On the right of the figure dot plots representing overall genomic synteny between the UViGs are indicated. OS3173, GBS41, JC39, and Conch37 represent TOSV, TGBSV, AJCSV, and ACSV, respectively.

**Figure 4.**
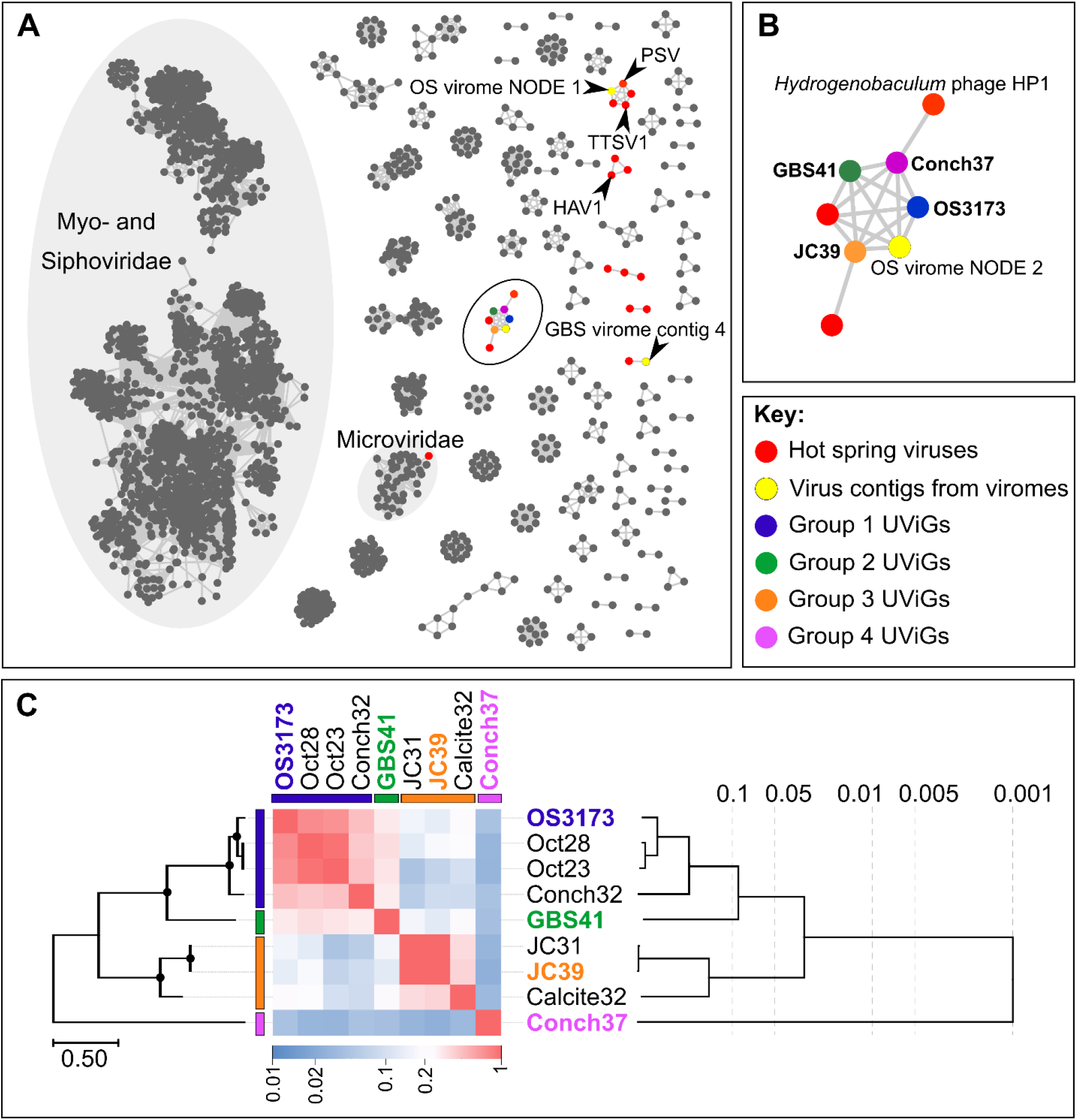
Relatedness inferred from gene-content between the OS3173-like UViGs. (A) Gene-sharing network inferred by vContact2 and visualized with Cytoscape 3.7.2. Nodes in the network represent cultivated or uncultivated viral genomes, while edges represent shared gene content between nodes. Viral contigs identified from hot spring microbial genomes are indicated in red, while contigs from either the Octopus Spring or Great Boiling Spring virus-enriched metagenomes are indicated in yellow (see Figure S4-7, File S1). The four representative UViGs are color-coded in their respective group colors. PSV, *Pyrobaculum* spherical virus; TTSV1, *Thermoproteus tenax* spherical virus 1; HAV1, Hyperthermophilic Archaeal virus 1. (B) Component of the gene-sharing network [circled in black on (A)] connecting the four representative UViGs together with the two outlier viruses, *Hydrogenobaculum* phage HP1 and another uncultivated virus from a pink streamer microbial community metagenome from Octopus Spring. One additional viral contig of the Octopus Spring virus-enriched metagenome was connected to the genus-level group Pyrovirus (OS virome NODE 2). (C) Genomic relatedness among the nine related UViGs based on normalized tBLASTx scores across the genomes (heatmap) with the PolA phylogeny depicted on left of the figure and a BioNJ phylogeny inferred from tBLASTx scores on the right. Bootstrap values above 95% on the polA phylogeny are indicated with circles at nodes. The phylogeny based on normalized tBLASTx scores of these UViGs, and their placement within the dsDNA viral reference sequences database, is indicated in Figure S3. OS3173, GBS41, JC39, and Conch37 represent TOSV, TGBSV, AJCSV, and ACSV, respectively.

For the classification of these nine UViGs, vContact2 was used to delineate genus-level groups for four representatives, one from each group in the Pol phylogeny, consisting of TOSV/OS3173 (Group 1), TGBSV (Group 2), *Aquificae* Joseph’s Coat Virus (AJCV) (Group 3), and *Aquificae* Conch Spring Virus (ACSV) (Group 4) (Figure 1; Table 2). The four representative UViGs were connected as a single component of the gene-sharing network (Figure 4A), with representatives from all four groups forming a single putative genus (proposed Pyrovirus) (Figure 4B). One outlier in the network, partially connected to the Pyrovirus component, was *Hydrogenobaculum* phage 1 (Figure 4A,B, S7) (Gudbergsdóttir et al., 2016), a 19,351 bp UViG recovered from a metagenome from Grensdalur, Iceland that was assigned to *Hydrogenobaculum* based on CRISPR spacer matches to genomes from cultivated *Hydrogenobaculum* strains. A second outlier (below the Pyrovirus group, Figure 4B) was obtained from a microbial metagenome of a pink streamer community from Octopus Spring. The gene-sharing network also illuminated some other viral contigs from the Octopus Spring and GBS viromes, belonging to gene-sharing sub-networks with PSV and *Thermoproteus tenax* spherical virus 1 (TTSV) (Ahn et al., 2006), Hyperthermophilic archaeal virus 1 (HAV) (Garrett et al., 2010), and *Microviridae*, among other isolated clusters. No genomes belonging to the primary *Myoviridae* or *Siphoviridae* networks were present in the hot spring metagenomes, reflecting the unique gene content of hyperthermophilic viruses.

**TABLE 2.** Summary of genomic features from four representative viral UViGs.

| UViG | Source | Length | %GC | Number<br>of genes | %<br>Coding | Proteins w/<br>function | % Proteins<br>w/ function |
| --- | --- | --- | --- | --- | --- | --- | --- |
| **TOSV**<br>**(OS3173)** | Octopus<br>Spring, WY | 37,265 | 37.1% | 49 | 95.1% | 21 | 35% |
| **TGBSV**<br>**(GBS41)** | Great Boiling<br>Spring, NV | 41,208 | 36.9% | 53 | 94.5% | 19 | 36% |
| **AJCSV**<br>**(JC39)** | Joseph's Coat<br>Spring, WY | 39,372 | 34.0% | 51 | 96.5% | 17 | 33% |
| **ACSV**<br>**(Conch37)** | Conch Spring,<br>WY | 37,258 | 35.5% | 50 | 94.8% | 15 | 30% |

### UNUSUAL BI-DIRECTIONAL GENOME REPLICATION STRATEGY AND UNIQUE GENOMIC FEATURES

The four representative genomes (TOSV/OS3173, TGBSV, AJCSV, ACSV) ranged from 37,256 bp to 41,208 bp in length, ranged in GC content from 34.0% to 37.1%, and encoded 48 to 53 open reading frames, with coding fraction ranging from 94.5% to 96.5% (Table 2, annotations found in supplemental File S2). The TOSV/OS3173 contig assembled into a circular genome, whereas the other genomes could not be circularized (Table 3) possibly due to lower coverage or incomplete assembly owing to population heterogeneity (Figure S2). For now, it is uncertain whether the genome is packaged as a circular molecule or whether it is packaged as a circularly permuted linear genome that circularizes only in the bacterial host. For all genomes, the transcriptional orientation of the ORFs is generally divided into a 23-26 kb set of contiguous genes on the same strand (clockwise in Figure 1), encoding 32-37 genes, and a smaller block on the other strand (counterclockwise in Figure 1), encoding 13-18 genes. As with most viral genomes, most genes are located in large blocks on the same strand. In each genome there are two to four instances of changes of strand involving one to two genes, except TGBSV, which consists exclusively of two large gene blocks, one on each strand. In the ACSV genome, there are two instances of a change of strand, each consisting of two genes. In each genome, a small (750 to 1,350 bp) intergenic region separated the sets of divergently transcribed genes, and this intergenic region also marked a strong divergence in GC skew. These features suggest bidirectional DNA replication beginning in the intergenic region around 36,429 bp of TOSV and the corresponding regions of the other viral genomes. These intergenic regions also contained repetitive elements predicted to form stem-loop structures, consistent with secondary structure typical of origins of replication. Many bacterial genomes are replicated bidirectionally, and their genomes have a G>C bias in the leading strand of replication and a C>G bias in the lagging strand (Képès et al., 2012); however, dsDNA phage do not typically replicate bidirectionally (Weigel and Seitz, 2006), and in this regard we suggest these viral genomes replicate more like mini bacterial genomes than typical phage genomes. Cultivation of one of the viruses would be necessary to test this hypothesis.

**TABLE 3.** Minimum Information about Uncultivated Virus Genomes (MIUViG) for the four representative UViGs.

| Metadata | TOSV<br>(OS3173) | TGBSV<br>(GBS41) | AJCSV<br>(JC39) | ACSV<br>(Conch37) |
| --- | --- | --- | --- | --- |
| **Source of UViG** | Viral fraction metagenome (virome) | Metagenome (not viral targeted) | Metagenome (not viral targeted) | Metagenome (not viral targeted) |
| **Sequencing approach** | Roche 454 | 454 GS FLX Titanium | Illumina HiSeq 2000, 2500 | Illumina HiSeq 2000, 2500 |
| **Assembly software** | CLC Genomics 8.0 (word size = 20, bubble size = 375), SPAdes v3.13.1 | SPAdes v 3.6.1 | SPAdes v 3.10.0 (--meta --only-assembler -k 21, 33, 55, 77, 99, 127) | SPAdes v 3.10.0 (--meta --only-assembler -k 21, 33, 55, 77, 99, 127) |
| **Viral identification software** | VirSorter, Earth's Virome pipeline, Inovirus detector pipeline | VirSorter, Earth's Virome pipeline, Inovirus detector pipeline | VirSorter, Earth's Virome pipeline, Inovirus detector pipeline | VirSorter, Earth's Virome pipeline, Inovirus detector pipeline |
| **Predicted genome type** | dsDNA | dsDNA | dsDNA | dsDNA |
| **Predicted genome structure** | Non-segmented | Non-segmented | Non-segmented | Non-segmented |
| **Detection type** | Independent sequence (UViG) | Independent sequence (UViG) | Independent sequence (UViG) | Independent sequence (UViG) |
| **Assembly quality** | Finished | High-quality draft | High-quality draft | High-quality draft |
| **Number of contigs** | 1 | 1 | 1 | 1 |

The presence of polymerase-, nuclease/recombinase-, and helicase-annotated genes in the smaller, counterclockwise set of genes in all four genomes suggests these genes might be transcribed earlier than the mainly structural genes in the larger, clockwise-facing block (Figure 1; Table S1, File S2). However, some genes encoding proteins associated with nucleotide metabolism were located among the clockwise-facing genes, including a DNA Pol III beta subunit (sliding clamp) in TOSV; a thymidylate kinase in TOSV, TGBSV, and AJCSV; and several genes that were found in only one of the four genomes, including site-specific DNA methylase (AJCSV), ribonucleotide reductase beta subunit (AJCSV), ATPase/kinase (AJCSV), and methyltransferases (ACSV). The location of these genes among the clockwise-facing part of the genomes and variability of these genes among the four UViGs suggest a variable and complex transcriptional/replication lifecycle for these viruses, or alternatively, that some nucleotide modification may be required during the lytic phase of infection.

Several genes encoding enzymes putatively involved in nucleic acid metabolism or DNA replication bear similarity to those in other viruses. ORF 3 of TOSV encodes a 119-amino acid protein with some similarity to a *Sulfolobus* virus DNA-binding protein that is highly conserved in diverse crenarchaeal viruses (Larson et al., 2007; Keller et al., 2007). TOSV and TGBSV both encode a putative sliding clamp beta subunit of DNA polymerase III, but they both lack an obvious clamp loader. Whether the viral replicase uses the host clamp loader or encodes an unrecognized clamp loader is unknown. Other viruses, including bacteriophage T4, encode sliding clamps, which have been shown to greatly increase processivity and the rate of replication (Trakselis et al., 2001). TOSV, TGBSV, and AJCSV each encode putative thymidylate kinases. Thymidylate kinases are encoded by a variety of viruses, including T4 and herpes simplex type 1 viruses. They are part of the nucleotide salvage pathway, typically have broad substrate activity, and are popular targets for antiviral drugs as they are often required for viability (Xie et al., 2019). ORF 5 in AJCSV encodes a putative site-specific DNA methylase. Viral genome methylation is a common epigenetic defense against host restriction-modification systems. Two putative methyltransferases of unknown activity are encoded by ORF 41 and ORF 42 of ACSV.

The counterclockwise-oriented genes included three major replicase-associated proteins that were conserved in all four UViGs: an ATP-dependent helicase (ORF 38 in TOSV), a nuclease/recombinase (ORF 37 in TOSV), and a large polyprotein encoding a Pol A with functionally active polymerase activity (OS3173 Pol) (ORF 36 in TOSV). The helicase genes contain two P-loop-containing nucleoside triphosphate hydrolase domains related to the DEAD-like helicase superfamily, but the similarity to functionally characterized orthologs is low. The Cas4-RecB-like nuclease (ORF 37 in OS3173) belongs to the PD-(D/E)XK nuclease superfamily, and may function as a single-stranded DNA-specific nuclease during replication and/or recombination, as these functions have been demonstrated for similar enzymes encoded by thermophilic archaeal viruses (Gardner et al., 2011; Guo et al., 2015).

ORF 36 in TOSV encodes a 1,606-amino acid polyprotein (OS3173 Pol), which was used to identify this group of viruses in the metagenomes (Figure 2). The amino-terminal region has conserved motifs that suggest primase and/or helicase function, including DUF927 (conserved domain with carboxy terminal P-loop NTPase) and COG5519 (Superfamily II helicases associated with DNA replication, recombination, and repair (Marchler-Bauer et al., 2011)). Consensus Walker A and Walker B motifs suggest NTP binding and hydrolysis likely associated with helicase activity (Walker et al., 1982). As reported previously (Schoenfeld et al., 2013), the viral *polA* genes are similar to the single genomic *polA* of *Aquificaceae* and *Hydrogenothermaceae*, as well as genes found as additional *polA* copies in a variety of other bacterial genomes, and to the nuclear-encoded, apicoplast-targeted DNA polymerases of several *Apicomplexa* species, typified by the Pfprex protein of *Plasmodium falciparum*. That enzyme is optimally active at 75°C (Seow et al., 2005), much higher than would be encountered during the *Plasmodium* life cycle, but similar to the optimal growth temperature of *Thermocrinis* and the geothermal springs sampled in this study, implying lateral gene transfer (Schoenfeld et al., 2013). Understanding the biochemical functions of the rest of the ORF 36 domains could reveal new thermostable accessory proteins for DNA amplification.

Most of the clockwise-facing genes that were annotated suggest these UViGs represent dsDNA tailed viruses belonging to the *Caudovirales*. Independent evidence that these viruses have dsDNA genomes comes from the initial study reporting the OS3173 PolA (Schoenfeld et al., 2008), because the viral DNA was amplified using a linker-dependent method that is specific for dsDNA. Furthermore, TOSV ORF 25, along with corresponding genes in the other UViGs, was annotated as a terminase large subunit, and ORF 24 was inferred to be a terminase small subunit, based on location immediately upstream of the large terminase, gene length (~300-400 bp), and a similar isoelectric point as other terminases. The terminase small subunit protein is a site-specific endonuclease that hydrolyzes viral DNA in preparation for packaging and encapsulation by the terminase large subunit (Kala et al., 2014). Terminase large subunit phylogeny has previously been used to infer the mechanism of packaging (Merrill et al., 2016, Chelikani et al., 2014); however, the terminases from this group of viruses was distant from those of well-studied viruses, so the mechanism of packaging could not be inferred (Figure S8). Immediately downstream of the putative terminase subunits in all genomes are two putative phage capsid proteins at ORF 26 and ORF 27 in TOSV. ORF 16 in TOSV was annotated as a portal protein, which forms dodecameric rings that play critical roles in virion assembly, DNA packaging, and DNA injection in *Caudovirales* (Prevelige and Cortines 2018). Additionally, TGBSV encodes a putative prohead protease (ORF 1), a WAIG tail domain protein (ORF 3), and a T7 tail fiber protein homolog (ORF 5), further supporting a relationship to *Caudovirales* and suggesting it encodes tail fibers typical of many *Caudovirales*. ORF 15 in TOSV was annotated as a lytic transglycosylase (lysin) based on the presence of a lysozyme-like domain. ORF 14 in TOSV was annotated as a holin based on the presence of three transmembrane domains, its small size (270 bp), and its location immediately upstream of ORF15. Also, the overlapping of open reading frames between ORFs 13, 14, and 15, suggests an anti-holin, holin, lysin operon, as found in numerous viruses. Together, these enzymes form the lysis cassette, which is common in *Caudovirales*, but not well understood in viruses of *Archaea* (Prangishvili 2013; Saier and Reddy 2015). There were also no lysogeny-related genes (e.g., integrases, excisionases or Cro/CI genes (Lima-Mendez et al., 2011, Shao et al., 2017) identified from these UViGs, suggesting a purely lytic lifestyle. As most of the clockwise-facing genes appear to be involved in viral packaging and lysis, these genes are predicted to be transcribed later than the counterclockwise-facing genes, as the lysis cassette is typically the last to be transcribed (Labrie et al., 2004, Young, 2014).

Each of the UViGs encode numerous hypothetical genes with no predicted function (~70%; including hits to known hypothetical proteins as well as those with no homology to known proteins), as is common in bacterial and archaeal viruses. Several of these were conserved among the genomes, but others were unique to each genome, or have diverged sufficiently that primary sequence conservation is difficult to discern. Many of the hypothetical proteins are related to genes found in different members of the *Aquificae*, consistent with the previous hypothesis that *Thermocrinis* and possibly other *Aquificae* are the putative hosts for these viruses.

### PUTATIVE HOSTS BELONG TO THE AQUIFICAE

Arrays of Clustered Regularly Interspaced Palindromic Repeats (CRISPRs) and related Cas (CRISPR associated) genes found in many bacterial and archaeal genomes (Grissa et al., 2007) provide a means to infer virus-host relationships (Gudbergsdóttir et al., 2016; Heidleberg et al., 2009; Snyder et al., 2010; Anderson et al., 2011; Roux et al., 2019a), as the CRISPR spacers provide a record of foreign nucleic acids that have been targeted by the CRISPR-Cas system. To determine the potential host range of these UViGs, genomes derived from isolates of *Hydrogenobaculum* sp. 3684, *Sulfurihydrogenibium yellowstonense* SS-5 ^T^, *Thermocrinis ruber* OC1/4^T^, and *Thermocrinis jamiesonii* GBS1^T^ were screened for CRISPR arrays with spacers matching the UViGs. *Hydrogenobaculum* sp. 3684 had six robust CRISPR clusters predicted, with the number of CRISPR spacers ranging between four and 50 in each cluster. In contrast, 19 CRISPR clusters were predicted for *Sulfurihydrogenibium yellowstonense* SS-5 ^T^, with the smallest having four spacer regions and the largest having 41. *T. ruber* OC1/4^T^ and *T. jamiesonii* GBS1^T^ genomes possessed six and four CRISPR clusters, ranging in the number of spacers between eight and 18, and four and 15, respectively. Each of these host genomes had one or more spacer with significant homology to the TOSV, TGBSV, and AJCSV genomes (Figure 5). No significant spacer matches were detected for ACSV. The six CRISPR spacer matches of the *T. ruber* OC1/4^T^ genome were somewhat distant (80-95% nucleic acid identity), which is reasonable considering that this organism was isolated from samples collected from Octopus Spring in 1994 (Huber et al., 1998), and the samples from which the UViGs were assembled were collected between 2007 and 2012. Furthermore, metagenomic studies of the pink streamer community in Octopus Spring revealed three dominant *Thermocrinis* populations, but each was distinct from *T. ruber* OC1/4^T^ (Takacs-Vesbach et al., 2013); thus, it is possible that the *T. ruber* OC1/4^T^ genotype is rarely encountered by TOSV. To assess this possibility, we analyzed *Thermocrinis* metagenome-assembled genomes (MAGs), as well as other MAGs from the *Aquificae*, from Octopus Spring (and other) metagenomes; however, the CRISPR arrays typically did not assemble with the respective MAGs, presumably because of non-native nucleotide word frequency associated with the foreign-derived CRISPR spacers (data not shown). Similarly, CRISPR spacer matches to the *Hydrogenobaculum* sp. 3684 and *Sulfurihydrogenibium yellowstonense* SS-5 ^T^ genomes were also distant (81-92%). By comparison, *T. jamiesonii* GBS1^T^, contained three arrays with four CRISPR spacers in total with significant identity to the TGBSV genome (>95%; ranging between 0 and 1 mismatch) (Figure 5B), providing strong evidence of the virus-host relationship.

**Figure 5.**
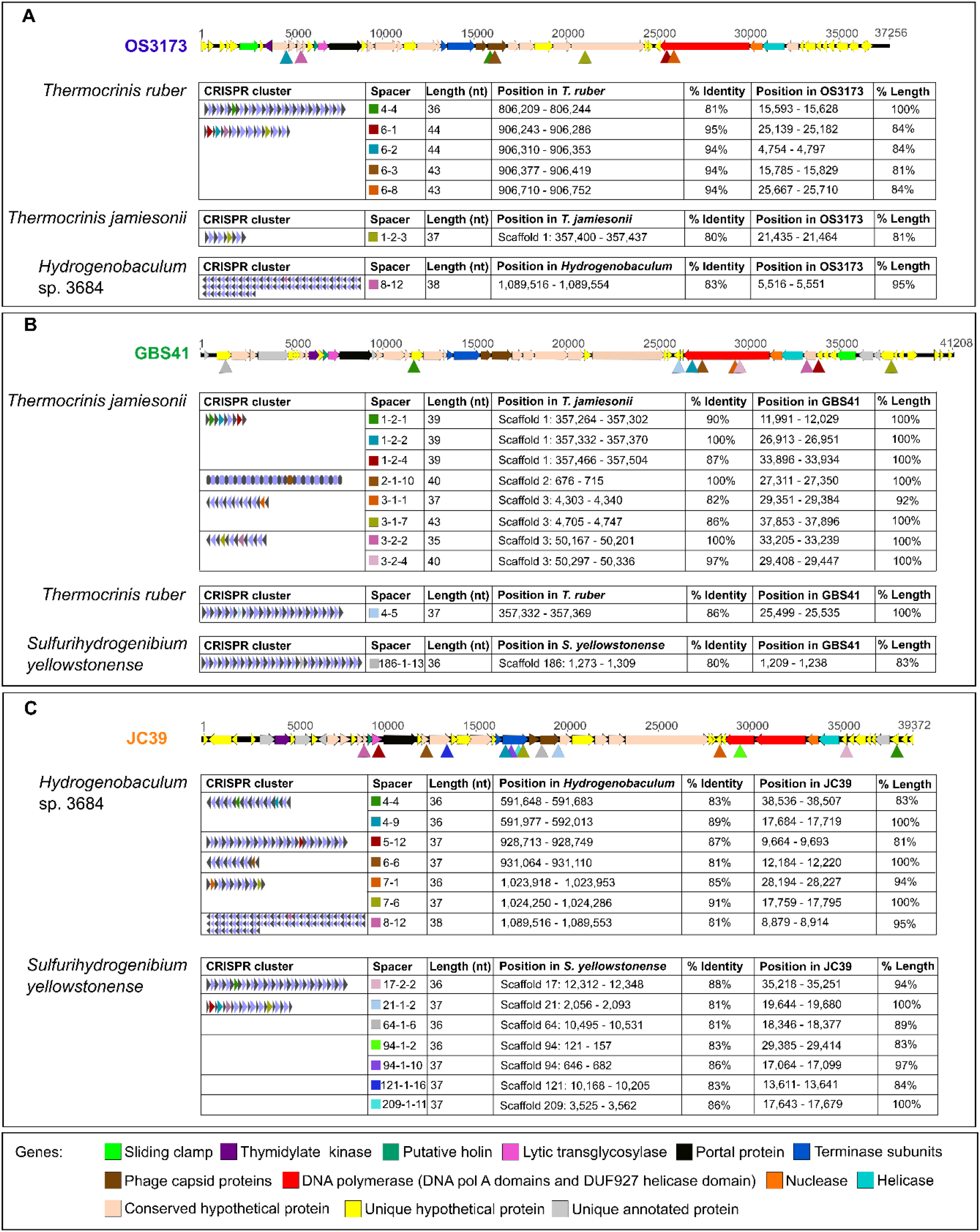
CRISPR spacer matches between viruses and *Aquificae* genomes. (A) Linearized map of the OS3173 genome with sites matching *Thermocrinis ruber* OC1/4^T^, *Thermocrinis jamiesonii* GBS1^T^, and *Hydrogenobaculum* sp. 3684 CRISPR spacer sequences denoted by triangles, and schematic and data on matching spacers. (B) Similar plot of the GBS41 genome with sites matching *Thermocrinis jamiesonii* GBS1^T^, *Thermocrinis ruber* OC1/4^T^, and *Sulfurihydorgenibium yellowstonense* SS-5^T^ CRISPR spacers. (C) Linearized map of the JC39 genome with corresponding CRISPR spacer sequence matches to *Hydrogenobaculum* sp. 3684 and *Sulfurihydorgenibium yellowstonense* SS-5^T^. OS3173, GBS41, and Conch37 represent TOSV, TGBSV, and ACSV, respectively.

The CRISPR spacers mapped to several different genes in the TOSV, TGBSV, and AJCSV genomes; however, the C-terminus of the PolA was targeted by spacers in each virus, and another two spacers mapped to the central portion of the PolA gene in TGBSV, suggesting that the C-terminus of the PolA is a functionally important antiviral target for the host (Figure 5). Accordingly, the C-terminal-encoding portion of the polA gene was among the most highly conserved regions of the genomes (Figure 3). The large capsid protein gene matched several spacers in TOSV and AJCSV, but not in TGBSV. The large terminase gene in AJCSV had matches to multiple CRISPR spacers, although this was not observed in the other two UViGs.

*Thermocrinis* is the dominant member of the pink streamer community in Octopus Spring (Reysenbach et al., 1994; Takacs-Vesbach et al., 2013) and the planktonic community in GBS (Cole et al., 2013); thus, it is reasonable to hypothesize that the natural host for the dominant viruses in these springs is *Thermocrinis*, as supported by shared gene content and CRISPR spacer matches. *Thermocrinis* is also extremely abundant in Little Hot Creek (Vick et al., 2010). Thus, we suggest that virus Groups 1 and 2, all encoding the larger polyprotein (Figure 2), associate with *Thermocrinis* as their putative host. These viruses are typified by TOSV (OS3173) and TGBSV, with the complete UViG of TOSV serving as the reference species for the group.

In contrast, *Sulfurihydrogenibium* was the dominant microorganism in Calcite Spring (Reysenbach et al., 2000) and Joseph’s Coat Spring was dominated by Archaea (Inskeep et al., 2013). We suggest that *Sulfurihydrogenibium* and/or *Hydrogenobaculum* are the most likely hosts for Group 3 and Group 4 viruses, especially as multiple hits were obtained to both these potential hosts with the AJCSV UViG. *Hydrogenobaculum* forms a distinct clade from *Thermocrinis*, *Hydrogenobacter*, *Aquifex*, and *Hydrogenivirga* within the *Aquificaceae*, and predominates in low pH springs (pH < 4.0) (Inskeep et al., 2013; Takacs-Vesbach et al., 2013). *Sulfurihydrogenibium* belongs to the sister family, *Hydrogenothermaceae*, and predominates in circumneutral springs (pH 6.5-7.8) (Takacs-Vesbach et al., 2013) and grows in a wide pH range in the lab (pH 5.0-8.8) (O’Neill et al., 2008). In this regard, it is noteworthy that some geothermal springs are poorly buffered and can change from circumneutral to highly acidic in both space and time, depending on the amounts and sources of geothermal and meteoric water that pool, and particularly on the source of sulfide, which can be oxidized to sulfuric acid by sulfide-oxidizing microorganisms (Nordstrom et al., 2009). Thus, it is possible that Group 3 and/or Group 4 viruses encounter and infect *Sulfurihydrogenibium* in circumneutral regions of the springs and *Hydrogenobaculum* in highly acidic regions, explaining the nearly equal numbers of CRISPR spacer matches to each organism. Additionally, the gene-sharing network and a neighbor-joining tree based on amino acid identity both suggested a distant relationship to *Hydrogenobaculum* phage 1 (Figure 3A,B, S3) (Gudbergsdóttir et al., 2016), a 19,351 bp UViG recovered from a metagenome from Grensdalur, Iceland that was assigned to *Hydrogenobaculum* based on CRISPR spacer matches to genomes from cultivated *Hydrogenobaculum* strains. Since the exact hosts of the Group 3 and Group 4 viruses are not conclusive, we suggest the names *Aquificae* Joseph’s Coat Virus (AJCV, high-quality draft genome) and *Aquificae* Conch Spring Virus (ACSV, high-quality draft genome) to represent the best genomes of Group 3 and Group 4.

### DESCRIPTION OF PROPOSED VIRUSES

(Py.ro.vi’rus. Gr. n. *pur,* fire; N.L. neut. n. Pyrovirus, “fire virus”, a thermophilic virus).

Based on the data presented here, we propose the following names and taxonomic relationships. Multiple genomic features suggest the nine novel UViGs belong to the order *Caudovirales*. The low overall sequence similarity and distinct placement of these taxa in gene-sharing networks suggest these viruses belong to an unclassified viral family and represent one putative genus-level group.

The proposed genus Pyrovirus accommodates TOSV (OS317), TGBSV, AJCV, and ACSV, with the complete genome of TOSV serving as the reference species for the genus. Members of this genus are predicted to infect *Aquificae* and are abundant in terrestrial geothermal springs. The estimated size of genomes in this genus range from 37 kb to 42 kb. The genomes contain genes encoding a thymidylate kinase, a holin, a lytic transglycosylase, a portal protein, large and small terminases, phage capsid proteins, DNA polymerase A (with fused or unfused DUF927 helicase domain), a nuclease and a helicase. Members of this genus are proposed to employ a complex bidirectional replication strategy.

## MATERIALS AND METHODS

### ISOLATION OF UNCULTURED VIRAL PARTICLES FROM OCTOPUS HOT SPRING AND GREAT BOILING SPRING

Virus particles were isolated from Octopus Hot Spring in Yellowstone National Park (Permit # YELL-2007-SCI-5240), Wyoming (N 44.5342, W 110.79812) in 2007 and from Great Boiling Spring (GBS), Nevada, (N 44.6614, W 119.36622) in October 2010, respectively. Temperature at the time and location of sampling was 87 °C at the outflow channel of Octopus Spring and 85 °C in the source pool of Great Boiling Spring.

For Octopus Spring samples, thermal water (between 200 and 630 liters) was filtered using a 100 kDa molecular weight cut-off (mwco) tangential flow filter (A/G Technology, Amersham Biosciences, GE Healthcare) and viruses and cells were concentrated to about 2 liters. The resulting concentrates were filtered through a 0.2 μm tangential flow filter to remove microbial cells. The viral fractions were further concentrated to about 100 mL using a 100 kDa tangential flow filter and 40 mL of viruses were further concentrated to 400 μL and transferred to SM buffer (0.1 M NaCl, 8 mM MgSO_4_, 50 mM Tris HCl, pH 7.5) by filtration in a 30 kDa mwco spin filter (Centricon, Millipore).

For the GBS viral sample tangential-flow filtration using a 30 kDa molecular weight cutoff Millipore Prep/Scale TFF-6 filter (catalog # CDUF006TT) was used to concentrate ~500 L of GBS water to ~2 L. Filtration was done in December 2010 with water from the GBS “A” site (Cole et al., 2013) with a temperature of 80-83 °C and pH of 7.15-7.2. The concentrated sample was stored on ice and transported to the laboratory, where it was pelleted by centrifugation at 4 °C for 10 minutes at 10,000 x g.

### ISOLATION OF VIRAL AND PLANKTONIC CELL DNA

*Serratia marcescens* endonuclease (Sigma, 10 U) was added to both viral preparations described above to remove non-encapsidated (non-viral) DNA. The reactions were incubated at 23°C for between 2 hours. EDTA (20 mM), sodium dodecyl sulfate (SDS) (0.5%) and Proteinase K (100 U) were added and the reactions were incubated at 56°C. Subsequently, sodium chloride (0.7M) and cetyltrimethylammonium bromide (CTAB) (1%) were added. The DNA was then extracted with chloroform, precipitated with isopropanol and washed with 70% ethanol. Yields of DNA ranged from 20 to 200 ng.

For preparation of cellular DNA from GBS, high molecular weight DNA was extracted from the pelleted cells essentially using the JGI bacterial DNA isolation CTAB protocol (https://jgi.doe.gov/user-programs/pmo-overview/protocols-sample-preparation-information/jgi-bacterial-dna-isolation-ctab-protocol-2012/). Briefly, this involved cell lysis with lysozyme (2.6 mg/mL), proteinase K (0.1 mg/mL), and SDS (0.5%), followed by purification of DNA by incubation with CTAB (1%) and sodium chloride (0.5 M), organic extraction, alcohol precipitation, treatment with RNase A (0.1 mg/mL), and an additional alcohol precipitation step.

### WHOLE-GENOME AMPLIFICATION OF VIRAL METAGENOMIC DNA

For the viral library that contained sequences of TOSV, a linker-based amplification method was used as described (Schoenfeld et al., 2008). For subsequent viral preparation isolated viral metagenomic DNA was amplified with an Illustra GenomiPhi V2 DNA amplification kit (G.E. Healthcare, Piscataway, NJ) following manufacturer’s protocol. Briefly, 9 μL sample buffer and 1 μL sample DNA were mixed and incubated at 95°C for 3 minutes and then placed on ice. Nine μL reaction buffer and 1 μL enzyme were then mixed and combined with the 10 μL sample and incubated for 2 hours at 30°C and 10 minutes at 65°C. The amplified DNA was then precipitated with NaCl and ethyl alcohol and resuspended in 40 μL water. The amplified DNA was debranched by adding 10 μL of 5X S1 nuclease buffer and 2 μL S1 nuclease (200 U; Thermo Fisher Scientific Inc., Waltham, MA), mixed and incubated at 25°C for 30 minutes and then 70°C for 10 minutes. The sample was reprecipitated twice with NaCl and ethyl alcohol and resuspended in 20 μL water. Several amplification reactions were prepared and used for DNA sequence analysis and to construct a large insert library in order to capture regions of the viral replisome.

### METAGENOMIC SEQUENCING AND ASSEMBLY

The amplified Octopus Spring viral metagenomic DNA was sequenced using Roche 454 chemistry at the Broad Institute (229,553 reads averaging 375 nucleotides each; 86,161,605 bases in total). The full read set was assembled *de novo* with CLC Genomics Workbench 8.0, using word size of 20 and bubble size of 375. A total of 5,143 contigs of length >500 were assembled with N50 = 1,818 bp, average length of 1,586 bp, maximum contig length of 35,614 bp (contig _4), and total assembly length of 8,156,404 bp. Of the 229,553 original reads, 66% (152,673 reads) were incorporated into contig assemblies >500 bp. Of the reads, 56.6% (86,379 reads) mapped to the largest contig (contig_4) at a stringency of 90%, which eventually was closed as Octopus Spring OS3173 virus (TOSV), resulting in an average coverage of 907-fold. The TOSV consensus viral sequence was finished by an iterative process of extending the ends of contig_4 with partially mapped reads until the extended consensus ends were found to overlap. This resulted in a 37,256 bp circular genome. A total of 99,924 reads were mapped to the finished genome (also at 90% stringency), and reads were found to map continuously across the joined overlap, consistent with a circular topology. Reads that did not map at 90% stringency were saved and remapped at relaxed stringency (80% identity over 80% length). These relaxed stringency reads were found to contain structural variants. The origin of the reported viral sequence was arbitrarily set to the beginning of the first ORF clockwise of the negative to positive GC skew transition (Figure 1). Viral contigs with lower coverage from the virus-enriched metagenome were obtained by reassembling the same reads using SPAdes v. 3.13.1 (Bankevich et al., 2012) with default parameters, except for the option “--only-assembler”.

Both cellular and amplified viral metagenomes from GBS were sequenced at the DOE Joint Genome Institute using Roche 454 GS FLX Titanium chemistry. Double-stranded genomic DNA samples were fragmented via sonication to fragments ranging between approximately 400 and 800 bp. These fragments were end-polished and ligated to Y-shape adaptors during 454 Rapid Library Construction. Clonal amplification of the library fragments was then performed in bulk through hybridization of the fragments to microparticle beads and subsequent emulsion-based PCR. Beads containing amplified DNA fragments were loaded into wells of a Pico Titer Plate (PTP) so that each well contained a single bead, followed by sequentially flowing sequencing reagents over the PTP. For the water-borne cell metagenome, a total of 355,082 reads were obtained ranging in length from 56 - 2,049 nucleotides producing 196,771,207 bases in total. During preprocessing through the DOE-JGI Metagenome Annotation Pipeline (MAP; https://img.jgi.doe.gov/m/doc/MetagenomeAnnotationSOP.pdf), 454 reads shorter than 150 bp and longer than 1,000 bp were removed. These reads were assembled with SPAdes v 3.6.1 (Bankevich et al., 2012), to a total of 315,164 contigs or sequences resulting in a total assembled size of 131,296,876 bases. Gene calling on the assembled sequences were done through the DOE-JGI MAP, resulting in the prediction of 271,395 RNA genes and 57,654 protein-coding genes. Through this pipeline, CRISPR array prediction was also done and a total of 508 CRISPR arrays were predicted to be present in the GBS cell metagenome. After binning with the DOE-JGI binning pipeline, a single *Thermocrinis jamiesonii* MAG was recovered. For the amplified viral metagenome or GBS virus-enriched metagenome, a total of 787,720 reads were sequenced ranging between 53 and 1,200 nucleotides for a total read library size of 392,631,172 bases. Read processing and assembly was also performed through the DOE-JGI MAP, in the same manner as the cellular metagenome. The virus-enriched metagenome had a total assembled size of 27,375,388 bases, which was divided over 55,185 contigs. In contrast to the cellular metagenome, only 137 RNA genes were predicted for this metagenome, supporting a low level of cellular contamination, and 74,087 protein-coding genes were predicted. A total of 60 CRISPR arrays were predicted.

### FUNCTIONAL ANNOTATION

ORFs in TOSV were identified by the GeneMarkS heuristic algorithm (Besemer et al., 2001). Open reading frames identified by GeneMarkS were submitted to NCBI BlastP (Altschul et al., 1990) using default settings for comparison with proteins in the public database.

Putative protein functions were inferred from searches against the NCBI nonredundant (nr) protein database with BLASTP (http://blast.ncbi.nlm.nih.gov), NCBI Conserved Domain Database (CDD) (http://ncbi.nlm.nih.gov/Structure/cdd) with CD-Search, UniProtKB with HMMer (http://hmmer.org), and CDD, Protein Data Bank (PDB), SCOPe 70 and Pfam with HHPred (https://toolkit.tuebingen.mpg.de/tools/hhpred). An E-value cutoff of 1e^−10^ was used for all tools. For each tool, the result with the lowest E-value that was not a “hypothetical protein” was chosen as the putative function predicted by that tool (Stamereilers et al., 2018). In some instances, putative function was assigned by synteny based on location and gene length (e.g., small terminase, holin).

In order to compile a composite annotation for all four of the UViGs used as representatives of the four PolA groups (i.e. Pyrovirus), all manual annotations were combined with functional annotations determined via the DOE-JGI MAP. Bidirectional BLASTp (Altschul et al., 1990) analyses were performed between all four viral sequences. Genes that were bidirectional best hits were considered homologous and robust annotations (separately identified as having the same function in at least two of the four UViGs) were transferred to all homologs. Where homologous genes had no functional annotation, or contradicting annotations between the reference sequences, the respective genes were denoted as encoding conserved hypothetical proteins.

### SINGLE-GENE TREES

In order to place the viral sequences identified to be close relatives of TOSV into phylogenetic context, two single-gene phylogenetic analyses were conducted on the protein sequences of firstly, the PolA from all viral scaffolds, together with the 3173 PolA-like sequences from Schoenfeld et al. (2013), and secondly, the large terminase subunit sequence. For the PolA phylogeny, the 3173 PolA-like sequences of *Thermocrinis* species were used for outgroup purposes based on previous studies (Schoenfeld et al., 2013). In contrast, the terminase phylogeny was unrooted, and reference sequences of Chelikani et al., (2014) were used to infer the potential packaging strategy of these viruses. Due to the variability present in these viral genes, the protein sequences were aligned based on structurally homologous protein domains with DASH (Rozewicki et al., 2019) in MAFFT v. 7 (Katoh et al., 2017; https://mafft.cbrc.jp/alignment/server/), with default settings. The appropriate protein model of evolution was determined for the respective alignments with ProtTest 3.4 (Darriba et al., 2011) and maximum likelihood analyses were conducted with RaxML v. 8.20 (Stamatakis, 2014). Branch support for the phylogenies was inferred from 1,000 bootstrap pseudoreplicates.

### PREDICTION OF PROTEIN DOMAINS

For the prediction of protein domains from the 3173 PolA-like sequences, a search of domain profiles based on hidden Markov Models was conducted through the EMBL-EBI hmmsearch tool (https://www.ebi.ac.uk/Tools/hmmer/search/hmmsearch) against the pfam database (El-Gebali et al., 2019). Protein family domains were predicted for all 3173 PolA-like protein sequences used in this study to determine whether the DUF 927 helicase and DNA pol A exo domains are fused to the pol A domain of the 3173 PolA-like proteins. Transmembrane domains for putative holins present in the four representative genomes from the proposed genus Pyrovirus were predicted through the TMHMM server (http://www.cbs.dtu.dk/services/TMHMM/).

### GENOME MAPS

Genome maps for the four reference sequences were constructed with CGView (Grant and Stothard, 2008; http://stothard.afns.ualberta.ca/cgview_server/). The GC content and skew for each genome was calculated with a step size of 1bp using a sliding window of 500bp. Protein-coding sequences were colored based on the homology inferences from the synteny analyses and the composite annotations for each genome. Breaks in the UViG sequences that were not circularized, i.e. TGBSV, AJCSV and ACSV, were indicated with red lines in all three tracks of the maps. The genome maps were rotated to align with that of TOSV for easier visualization.

### RELATIVE ABUNDANCE OF VIRAL CONTIGS IN VIROMES

From the metagenomes analyzed, viral genomes were predicted with VirSorter v. 1.0.5 (Roux et al., 2015), Earth’s Virome pipeline (Paez-Espino et al., 2016) and Inovirus detector pipeline v. 1.0 (https://bitbucket.org/srouxjgi/inovirus/src/master/) (Roux et al., 2019b). From the respective viral-enriched metagenomes, 372 contigs were obtained with 42 contigs ≥10,000 bp. Dereplication was done with an Average Nucleotide Identity (ANI) of 95% over an alignment fraction of 85% to obtain 320 non-redundant contigs. Contig coverage was estimated by mapping reads from individual metagenomes to the 320 non-redundant viral contigs using BBMap v. 38.67 (https://www.osti.gov/biblio/1241166-bbmap-fast-accurate-splice-aware-aligner). Only reads that mapped at ≥95% nucleotide identity were considered and contig coverage was set at 0 if less than 70% of the contig’s length was covered by metagenomic reads, or as the average read depth per position otherwise, as typical for UViG analysis (Roux et al., 2019a).

### VIRAL CLASSIFICATION

All contigs ≥10,000 bp obtained from the virus-enriched metagenomes, together with the four representative UViGs were used as input with the viral reference sequence database (RefSeq v94), to automatically delineate genus-level groups based on shared gene content in vContact2 using default parameters (Bin Jang et al., 2019). The resulting gene-sharing network was viewed and edited in Cytoscape 3.7.2 (http://cytoscape.org), using a prefuse force directed layout.

### PROTEOMIC TREE AND SYNTENY ANALYSES

In order to confirm the relationships among the nine UViGs, a proteomic tree was constructed with ViPtree (Nishimura et al., 2017; https://www.genome.jp/viptree/). This Neighbor-Joining (NJ) tree is constructed by computing genome-wide tBLASTx similarity scores (McGinnis and Madden, 2004) among all submitted and all reference viral sequences. These similarity scores were then used to construct a distance matrix used for constructing a BIONJ tree. Based on previous results, the nucleic acid type was specified as dsDNA, with prokaryotes indicated as the potential hosts. Gene predictions as performed above were used for the UViGs. This process was repeated for the 10 UViGs with the highest coverage in the two virus-enriched metagenomes (i.e. from Octopus Spring and Great Boiling Spring). For depicting synteny, the genome alignments based on tBLASTx analyses, as inferred with ViPtree, was used.

### HOST IDENTIFICATION FOR ABUNDANT VIRUSES IN GREAT BOILING SPRING AND OCTOPUS SPRING

The ten viruses with the highest coverage in Great Boiling Spring and Octopus Spring respectively, were identified from the viral metagenomes. In order to identify potential hosts for these viruses, a two-pronged approach was employed. The first approach consisted of identifying potential prophages in bacterial and archaeal genomes, while the second approach consisted of identifying CRISPR spacers in host genomes matching the viral sequences.

For the identification of potential prophages matching the viral sequences, BLASTn analyses were conducted with the 10 viruses with the highest coverage in each spring to the DOE JGI/IMG isolate genome database (Chen et al., 2019), as well as the NCBI Whole Genome Shotgun (WGS) and RefSeq Genomic (refseq_genomic) databases.

For the second approach, CRISPR clusters were used from all metagenomes, SAGs and isolate genomes, available on IMG for Octopus Spring and Great Boiling Spring. All CRISPR spacer regions available on IMG for these genomes were used for further analysis. Those single-amplified genomes and isolate genomes that did not have CRISPR prediction results available on IMG were analyzed with CRISPRCasFinder (https://crisprcas.i2bc.paris-saclay.fr/CrisprCasFinder/Index; Couvin et al., 2018). All predicted spacer regions were then compared to the ten most covered virus sequences in each spring using BLASTn (BLAST v.2.2.31; Altschul et al., 1990) with custom settings (-word_size 7 -gapopen 10 - gapextend 2 -penalty -1 -outfmt 6 -dust no). For the spacer comparisons from the metagenomes, only spacer regions with matches over 100% of the length of the spacer were considered, while matches over 80% of the length of the spacers were considered for SAGs and isolate genomes. Resulting BLAST hits were then further limited to those with a percentage identity of ≥80% and an Expect(e)-value of ≤0.00001.

For the CRISPR spacer detection of the four representative UViGs to *Hydrogenobaculum* sp. 3684, *Sulfurihydrogenibium yellowstonense* SS-5 ^T^, *Thermocrinis ruber* OC1/4^T^, and *Thermocrinis jamiesonii* GBS1^T^, these microbial isolate genomes were subjected to CRISPR array prediction with CRISPRCasFinder. The resulting CRISPR arrays with a confidence level of three or above were further analyzed. All predicted spacer sequences were subjected to BLASTn analyses against TOSV, TGBSV, AJCSV and ACSV as described above.

### RECRUITMENT PLOTS

To visualize the level of variability within the viral populations and coverage across the UViGs for Octopus Spring and Great Boiling Spring, raw sequence reads were recruited to the UViGs of TOSV and TGBSV. The UViGs were used to construct BLAST databases using makeblastdb in BLAST v. 2.2.31. Following this, BLASTn analyses were conducted with each UViG database as reference and their respective metagenomic reads from which they were assembled, as query. Default settings for BLAST analyses were used apart from specifying tabular format for the data output (-outfmt 6), reporting a single HSP per subject sequence (-max_hsps 1) and keeping a single alignment per subject sequence (-max_target_seqs 1). The BLAST results were formatted with BlastTab.catsbj.pl (http://enveomics.blogspot.com/2013/01/blasttabcatsbjpl.html) limiting the identity of hits to report to 30%, and these data was then subjected to recruitment plot construction with enve.recplot2 in the Enveomics Collection (https://github.com/lmrodriguezr/enveomics; Rodriguez-R & Konstantinidis, 2016) in RStudio v. 3.6.1. To compare obtained recruitment plots to the genomic architecture of the UViGs, annotated UViGs were visualized with Geneious R7 (Biomatters) and edited in Inkscape v. 0.92.

### SEQUENCE ACCESSION NUMBERS

The individual sequence reads from the 2007 Octopus hot spring viral sample can be accessed at http://data.imicrobe.us/search?query=great+boiling+spring. The quality-filtered reads is being submitted to the National Center for Biotechnology Information (NCBI) Sequence Read Archive (SRA). The other accession numbers for the eight TOSV relatives can be found in the DOE-JGI IMG/M (Chen et al., 2019) website (http://img.jgi.doe.gov/m) under IMG Scaffold ID numbers found in Table 4. The four representative UViGs (TOSV, TGBSV, AJCV and ACSV) are also being submitted to the NCBI (https://www.ncbi.nlm.nih.gov/) under the nucleotide database.

## ACKNOWLEDGEMENTS

We thank the Gordon and Betty Moore Foundation for funding the sequence of the viral metagenome from Octopus Spring, and Matt Henn at the Broad Institute for 454 sequencing. This research was supported by the United States National Science Foundation grant DEB 1557042, United States Department of Energy grant DE-EE-0000716, and the Joint Genome Institute at the DOE (CSP-182). The work conducted by the U.S. Department of Energy Joint Genome Institute, a DOE Office of Science User Facility, is supported by the Office of Science of the U.S. Department of Energy under Contract No. DE-AC02-05CH11231.

