## Supplementary Figures and Tables for "Diversity and Distribution of a Novel Genus of Hyperthermophilic *Aquificae* Viruses Encoding a Proof-reading Family-A DNA Polymerase"

**Table S1. Minimum Information about additional Uncultivated Virus Genomes (MIUViG).**

| Metadata | Oct28 | Oct23 | Conch32 | JC31 | Calcite32 |
|---|---|---|---|---|---|
| **Source of UViG** | Metagenome (not viral targeted) | Metagenome (not viral targeted) | Metagenome (not viral targeted) | Metagenome (not viral targeted) | Metagenome (not viral targeted) |
| **Sequencing approach** | Illumina HiSeq 2000, Illumina HiSeq 2500 | Illumina HiSeq 2000, Illumina HiSeq 2500 | Illumina HiSeq 2000 | Illumina HiSeq 2000 | Sanger |
| **Assembly software** | SPAdes v 3.10.0 (--meta --only-assembler -k 21, 33, 55, 77, 99, 127) | SPAdes v 3.10.0 (--meta --only-assembler -k 21, 33, 55, 77, 99, 127) | SPAdes v 3.10.0 (--meta --only-assembler -k 21, 33, 55, 77, 99, 127) | SPAdes v 3.10.0 (--meta --only-assembler -k 21, 33, 55, 77, 99, 127) | Phrap |
| **Viral identification software** | Viral polA BLAST | Viral polA BLAST | Viral polA BLAST | Viral polA BLAST | Viral polA BLAST |
| **Predicted genome type** | dsDNA | dsDNA | dsDNA | dsDNA | dsDNA |
| **Predicted genome structure** | Non-segmented | Non-segmented | Non-segmented | Non-segmented | Non-segmented |
| **Detection type** | Independent sequence (UViG) | Independent sequence (UViG) | Independent sequence (UViG) | Independent sequence (UViG) | Independent sequence (UViG) |
| **Assembly quality** | Genome fragment | Genome fragment | Genome fragment | Genome fragment | Genome fragment |
| **Number of contigs** | 1 | 1 | 1 | | 1 |

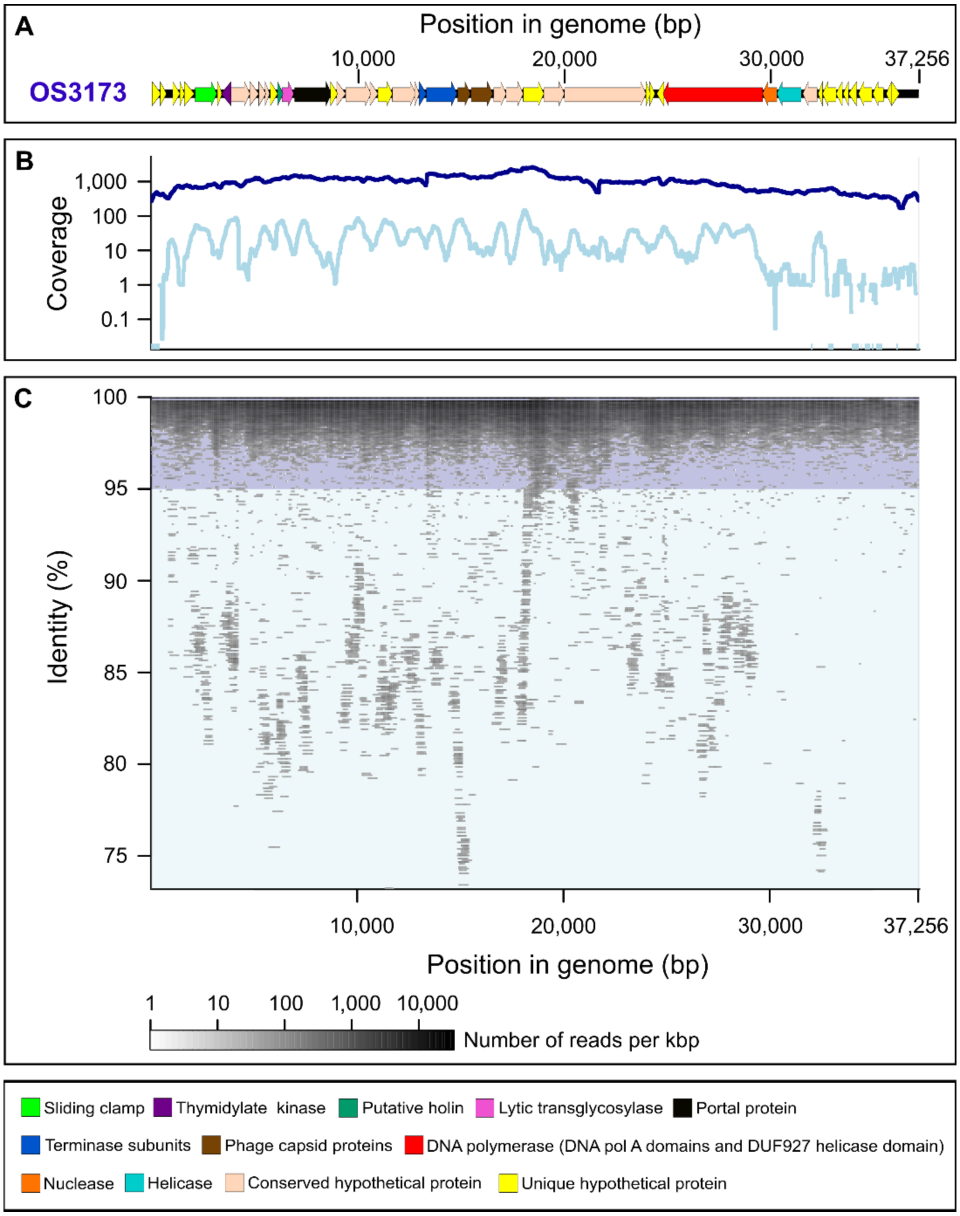


**Figure S1. Recruitment of the OS3173 virus genome.** (A) Linearized map of the OS3173 genome. Arrows denote putative direction of transcription. Genes are color coded as shown in the bottom panel. (B) Coverage across the genome at 95% nucleotide identity (dark blue) and between 95% and 80% nucleotide identity (light blue). (C) Plot of individual reads across the genome. Shades of blue correspond to coverage shown in panel B.

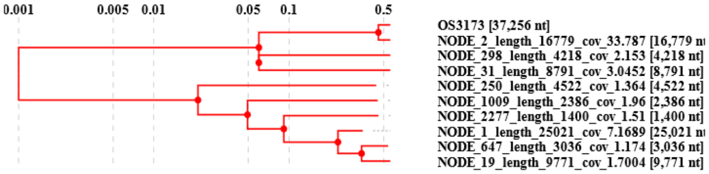


Normalized tBLASTx Score

| | OS3173 | NODE_2 | NODE_298 | NODE_31 | NODE_250 | NODE_1009 | NODE_2277 | NODE_1 | NODE_647 | NODE_19 |
|---|---|---|---|---|---|---|---|---|---|---|
| OS3173 | 1 | 0.7951 | 0 | 0 | 0 | 0 | 0 | 0 | 0 | 0 |
| NODE_2 | 0.7951 | 1 | 0 | 0 | 0 | 0 | 0 | 0 | 0 | 0 |
| NODE_298 | 0 | 0 | 1 | 0 | 0 | 0 | 0 | 0 | 0 | 0 |
| NODE_31 | 0 | 0 | 0 | 1 | 0 | 0 | 0 | 0 | 0 | 0 |
| NODE_250 | 0 | 0 | 0 | 0 | 1 | 0.1152 | 0.1059 | 0.3545 | 0 | 0 |
| NODE_1009 | 0 | 0 | 0 | 0 | 0.1152 | 1 | 0.173 | 0.3647 | 0 | 0 |
| NODE_2277 | 0 | 0 | 0 | 0 | 0.1059 | 0.173 | 1 | 0.4721 | 0 | 0 |
| NODE_1 | 0 | 0 | 0 | 0 | 0.3545 | 0.3647 | 0.4721 | 1 | 0.6279 | 0.4847 |
| NODE_647 | 0 | 0 | 0 | 0 | 0 | 0 | 0 | 0.6279 | 1 | 0.5878 |
| NODE_19 | 0 | 0 | 0 | 0 | 0 | 0 | 0 | 0.4847 | 0.5878 | 1 |

**Figure S2.** Similarity matrix inferred from normalized tBLASTx scores and associated neighbor-joining tree of ten viral contigs with the highest coverage from the Octopus Spring virus-enriched metagenome. Contig length and sequence identifiers are noted on the labels on the distance tree. A phylogenetic analysis relating these contigs to known dsDNA viral genomes is shown in Figure S4. Overall, the ten contigs with the highest coverage obtained from the Octopus Spring virus-enriched metagenome were grouped into at least four distinct clusters based on the viral proteomic tree approach. The first cluster contains OS3173 (TOSV) together with contig NODE 2 (also visible in Figure 4, members of Pyrovirus), two singleton clusters consisting of NODE 298 and NODE 31, respectively, and a large cluster containing NODE 1 (see Figure 4), NODE 250, NODE 1009, NODE 2277, NODE 647 and NODE 19 related to *Pyrobaculum* spherical virus and *Thermoproteus tenax* spherical virus 1 (Figure 3).

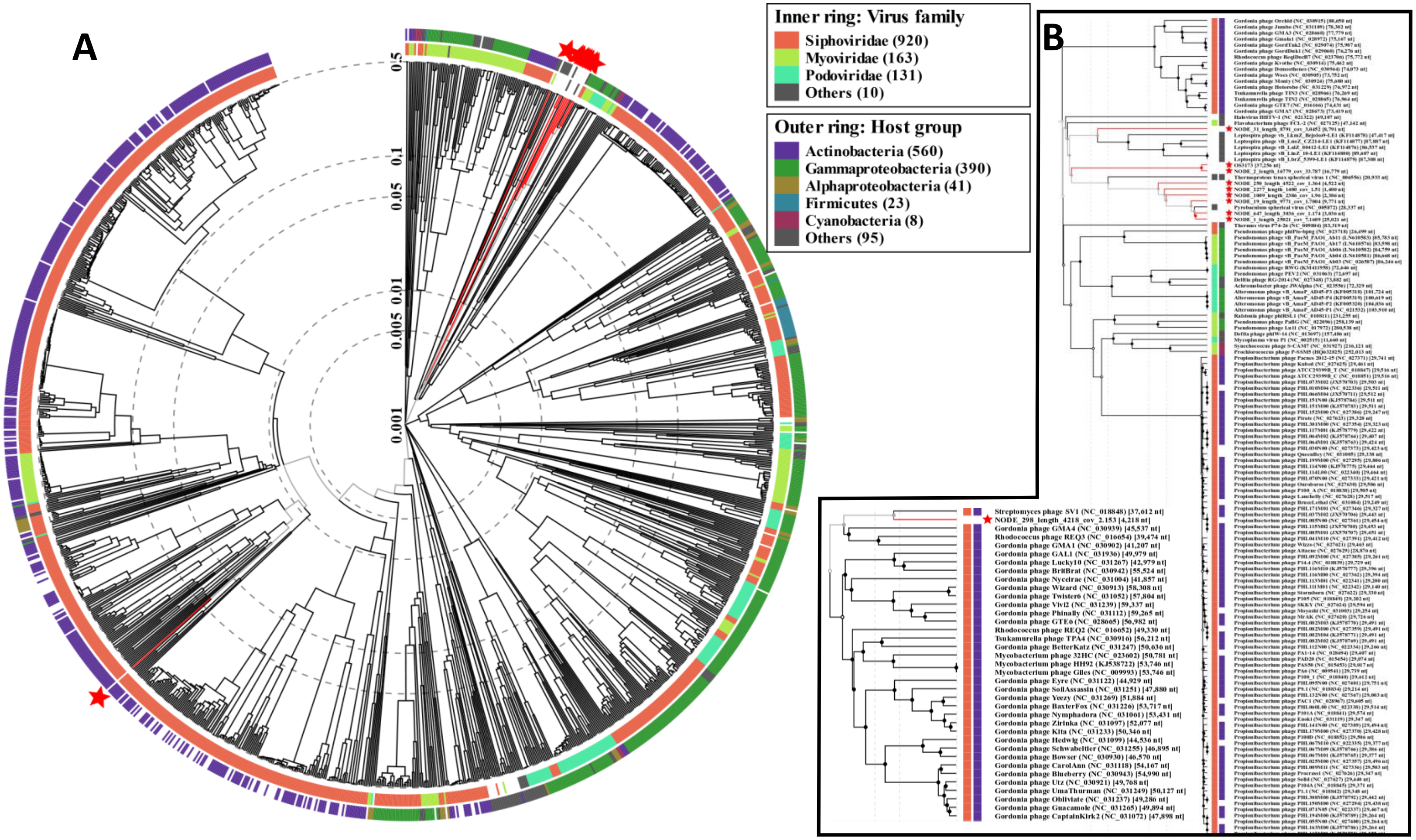


**Figure S3.** (A) Neighbor-joining tree of ten viral contigs with the highest coverage from Octopus Spring virus-enriched metagenome within the context of other dsDNA viral genomes. The placement of the ten contigs with the highest coverage from the Octopus Spring virus-enriched metagenome is indicated with red stars. (B) Subtrees containing these 10 viral contigs, indicated in red, with their closest relatives. The same clusters were obtained as from results of the gene-sharing network together with the tBLASTx relationships. The first cluster contains OS3173(TOSV) together with contig NODE 2 (also visible in Figure 4, members of Pyrovirus), and groups as sister to the large cluster containing NODE 1 (see Figure 4, S3), NODE 250, NODE 1009, NODE 2277, NODE 647, NODE 19, *Pyrobaculum* spherical virus and *Thermoproteus tenax* spherical virus 1 (Figure 4, S3). The two singletons were grouped with *Leptospira* and *Streptomyces* phages, respectively.

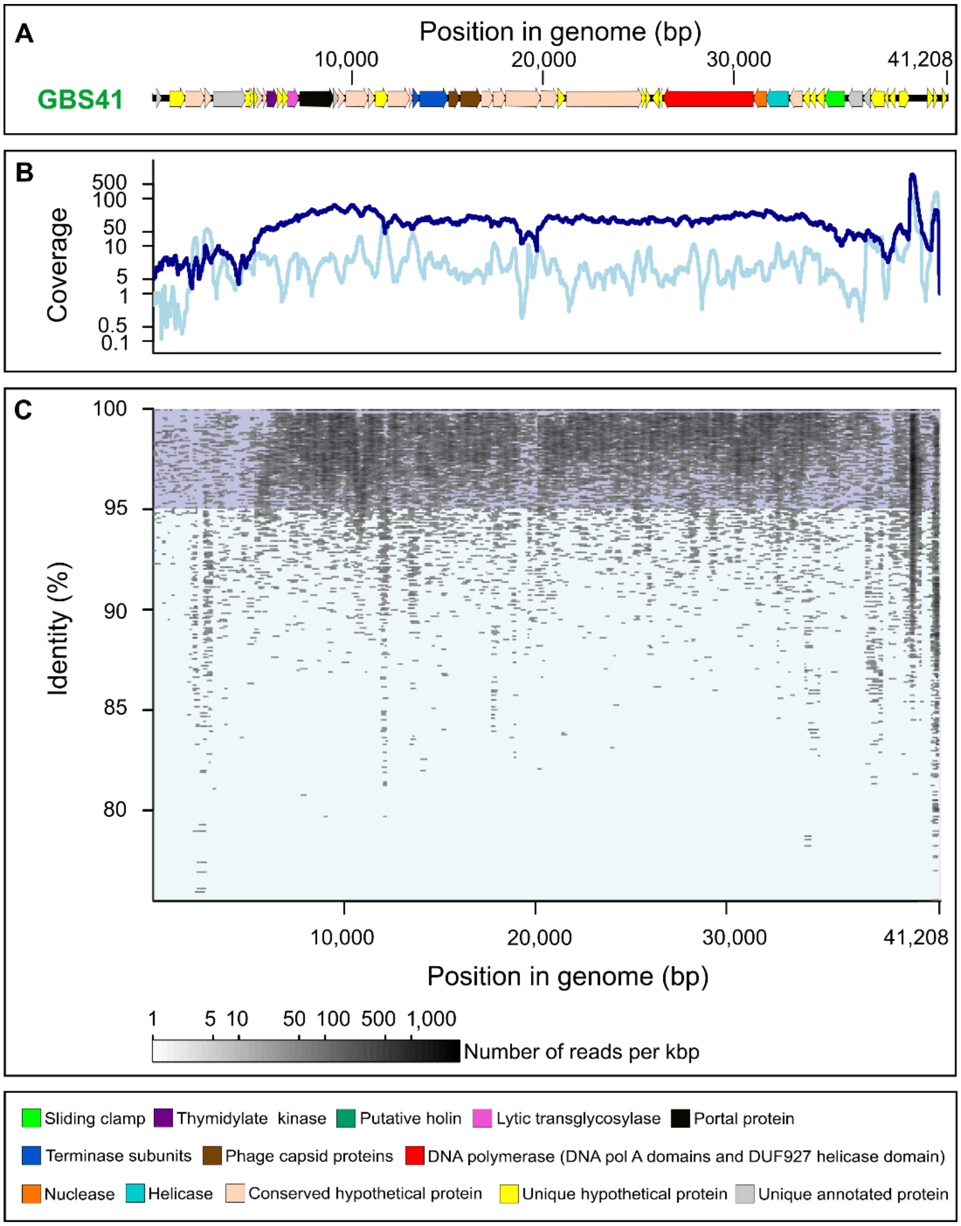


**Figure S4. Recruitment of the GBS41 virus genome.** (A) Linearized map of the OS3173 genome. Arrows denote putative direction of transcription. Genes are color coded as shown in the bottom panel. (B) Coverage across the genome at 95% nucleotide identity (dark blue) and between 95% and 80% nucleotide identity (light blue). (C) Plot of individual reads across the genome. Shades of blue correspond to coverage shown in panel B.

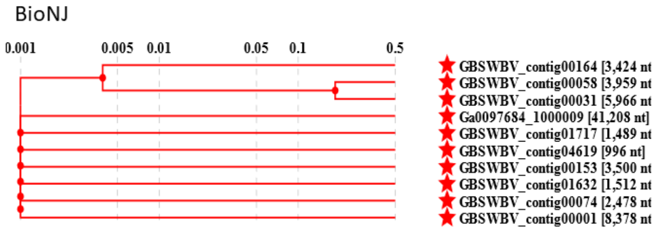


10 Most abundant viruses GBS - Related viruses

Normalized tBLASTx Score

| | contig00164 | contig00058 | contig00031 | Ga0097684 | contig01717 | contig04619 | contig00153 | contig01632 | contig00074 | contig00001 |
|---|---|---|---|---|---|---|---|---|---|---|
| GBSWBV_contig00164 | 1 | 0.0178 | 0 | 0 | 0 | 0 | 0 | 0 | 0 | 0 |
| GBSWBV_contig00058 | 0.0178 | 1 | 0.3706 | 0 | 0 | 0 | 0 | 0 | 0 | 0 |
| GBSWBV_contig00031 | 0 | 0.3706 | 1 | 0 | 0 | 0 | 0 | 0 | 0 | 0 |
| Ga0097684_1000009 | 0 | 0 | 0 | 1 | 0 | 0 | 0 | 0 | 0 | 0 |
| GBSWBV_contig01717 | 0 | 0 | 0 | 0 | 1 | 0 | 0 | 0 | 0 | 0 |
| GBSWBV_contig04619 | 0 | 0 | 0 | 0 | 0 | 1 | 0 | 0 | 0 | 0 |
| GBSWBV_contig00153 | 0 | 0 | 0 | 0 | 0 | 0 | 1 | 0 | 0 | 0 |
| GBSWBV_contig01632 | 0 | 0 | 0 | 0 | 0 | 0 | 0 | 1 | 0 | 0 |
| GBSWBV_contig00074 | 0 | 0 | 0 | 0 | 0 | 0 | 0 | 0 | 1 | 0 |
| GBSWBV_contig00001 | 0 | 0 | 0 | 0 | 0 | 0 | 0 | 0 | 0 | 1 |

**Figure S5.** Distance matrix and neighbor-joining tree of ten viral contigs with the highest coverage from the Great Boiling Spring viral metagenome. Contig length and sequence identifiers are noted on the labels on the distance tree with sequence identifiers deposited in the DOE-JGI IMG/M. A Neighbor-Joining phylogenetic analysis relating these contigs to known dsDNA viral genomes is shown in Figure S6. The ten contigs with the highest coverage obtained from the Great Boiling Spring virus-enriched metagenome were grouped into eight distinct clusters based on the viral proteomic tree approach. The first cluster contains contig00164, contig00058 and contig00031, with GBS41 (TGBSV, Ga0097684_1000009) as the sole member of Pyrovirus, while all other contigs showed no similarity among them based on normalized tBLASTx scores.

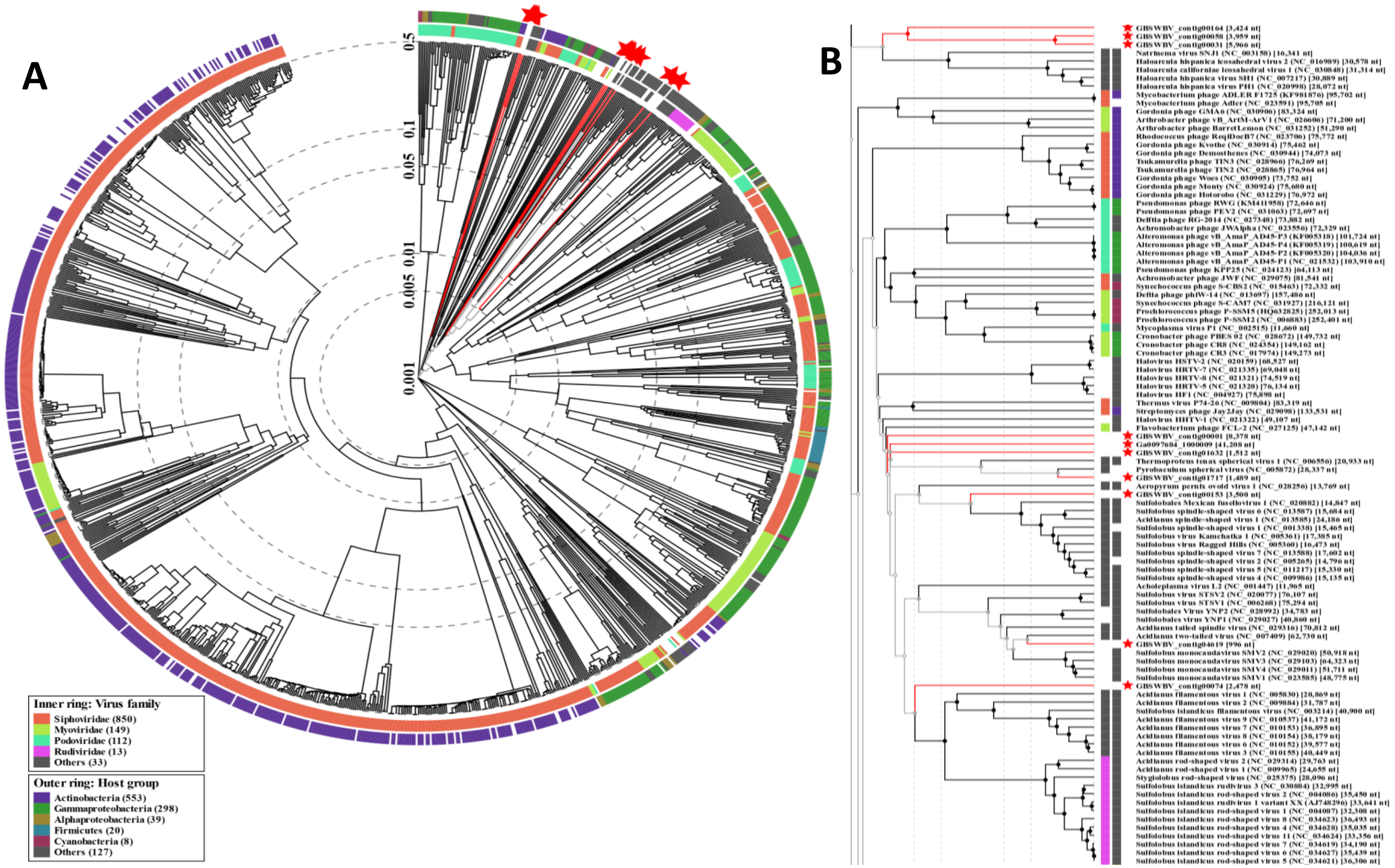


**Figure S6.** (A) Neighbor-joining tree of the ten viral contigs with the highest coverage from the Great Boiling Spring (GBS) viral metagenome within the context of other dsDNA viral genomes. (B) Subtree containing these 10 GBS viral contigs indicated in red, with their closest relatives. A single grouping of contig00164, contig00058 and contig00031 was placed on a distinct branch as sister to *Natrinema* and *Haloarcula* viruses, contig01717 were placed as sister to *Pyrobaculum* spherical virus, three contigs were placed as somewhat related to *Sulfolobales* and *Sulfolobus* viruses, while the placement of the remaining contigs were uncertain. However, the placement of all of these viral contigs, with the exception of GBS41, are not reliable due to the small size of these contigs.

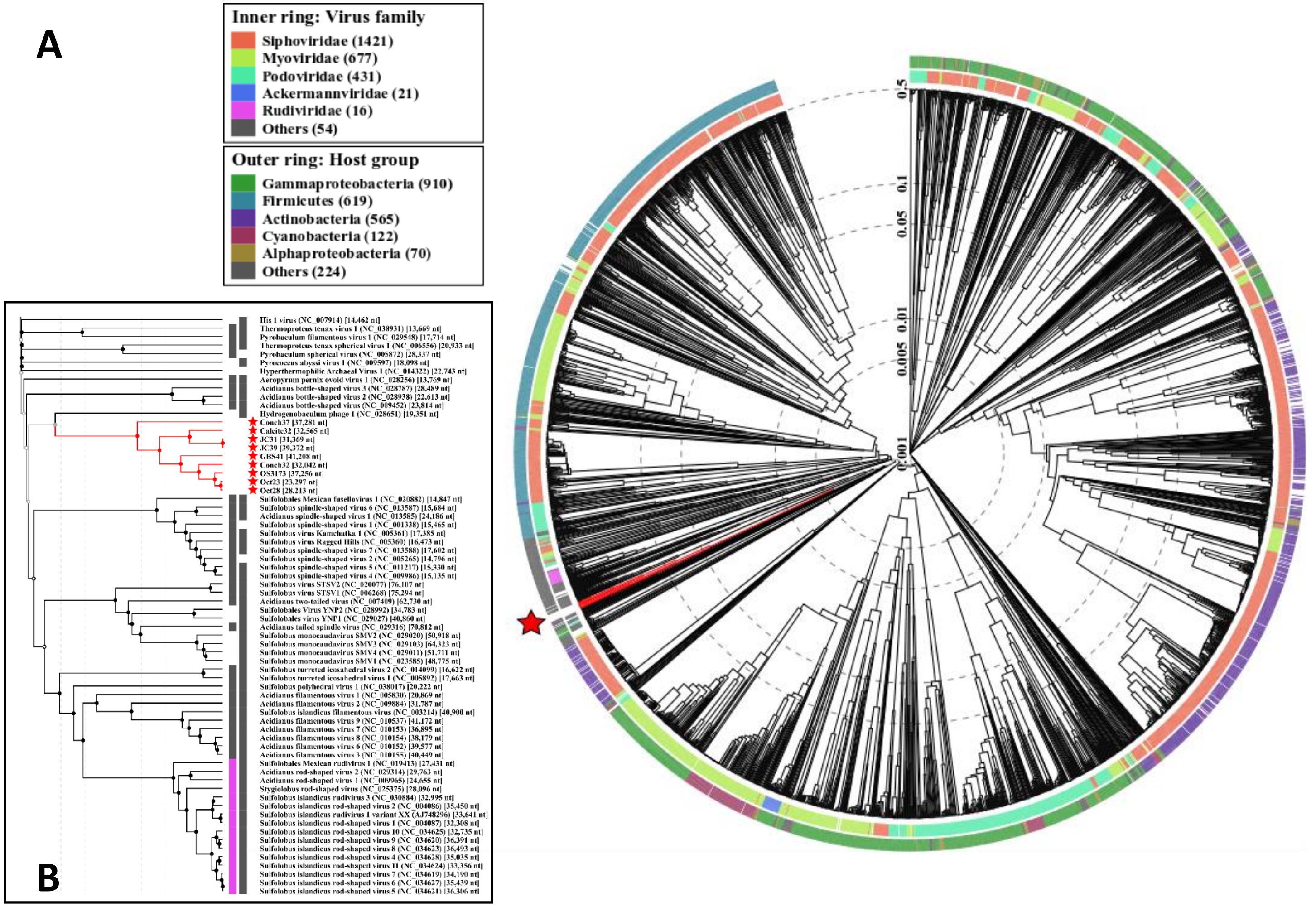


**Figure S7.** Relationships between viral contigs are inferred from normalized tBLASTx scores using the viral proteomic tree approach. (A) Neighbor-joining tree of the OS3173-like UViGs within the context of other dsDNA viral genomes. The red star denotes the placement of members of the putative novel genus Pyrovirus. (B) Subtree containing the OS3173-like UViGs and closest related dsDNA viral genomes. Of all available dsDNA viral reference sequences, only *Hydrogenobaculum* phage HP1 showed any similarity to members of the putative genus Pyrovirus, with a very low normalized tBLASTx score of 0.02 to Conch37.

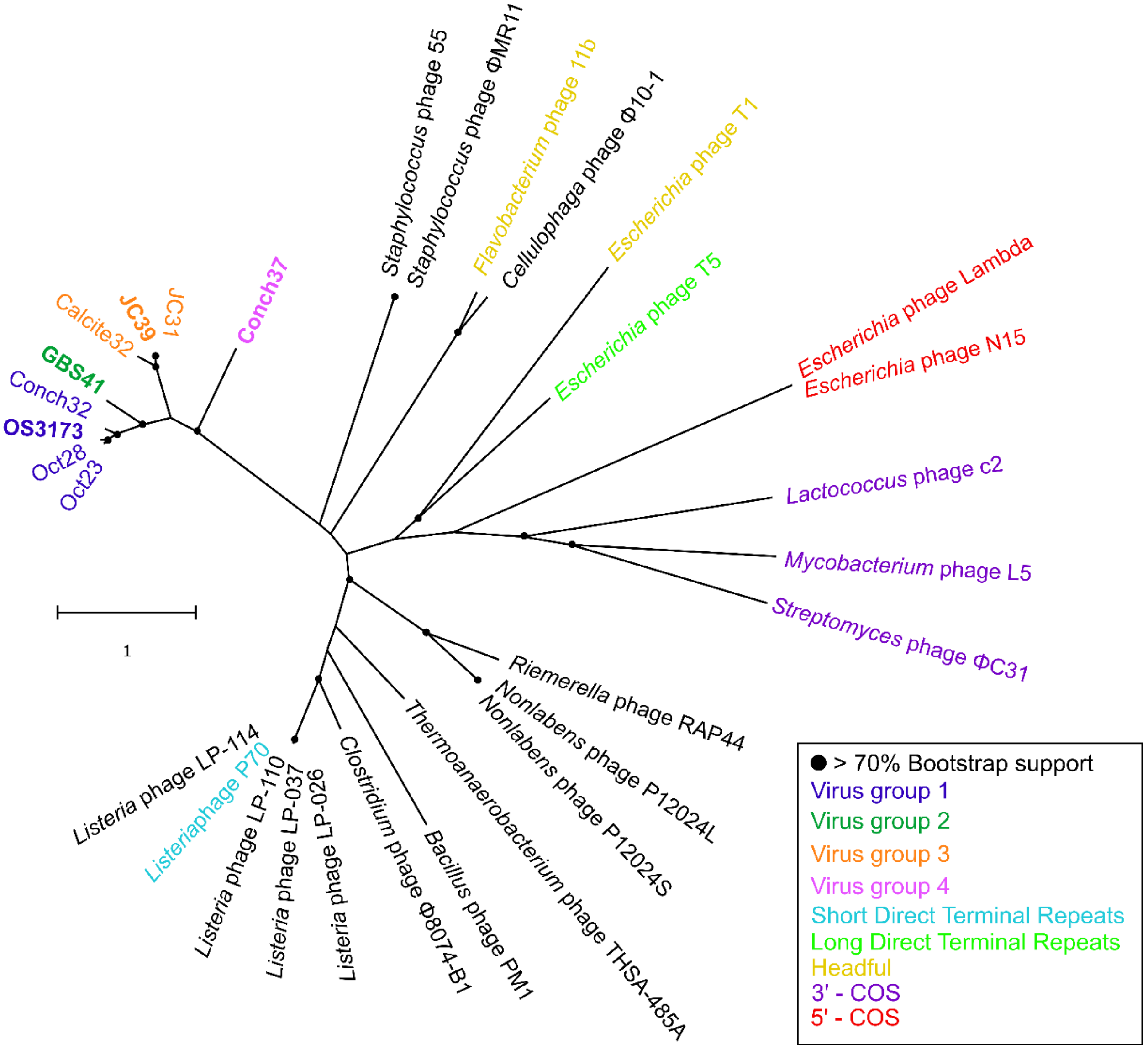


**Figure S8.** Maximum-likelihood tree of the protein sequences of large terminases. Branch support was inferred from 1,000 bootstrap pseudoreplicates. Reference sequences from Chelikani et al., 2014 was used.
